# Neurobehavioral precursors of compulsive cocaine-seeking in dual fronto-striatal circuits

**DOI:** 10.1101/2022.11.09.515779

**Authors:** Jolyon A. Jones, Aude Belin-Rauscent, Bianca Jupp, Maxime Fouyssac, Stephen J. Sawiak, Katharina Zuhlsdorff, Peter Zhukovsky, Lara Hebdon, Clara Velazquez Sanchez, Trevor W. Robbins, Barry J. Everitt, David Belin, Jeffrey W. Dalley

## Abstract

Only some individuals using drugs recreationally eventually become addicted, and persist in drug seeking and taking despite adverse consequences. The neurobehavioral determinants of this individual vulnerability have not been fully elucidated. We report that in drug naïve rats the future tendency to develop compulsive cocaine seeking is characterised by behavioral stickiness-related functional hypoconnectivity between the prefrontal cortex and posterior dorsomedial striatum in combination with impulsivity-related structural alterations in the infralimbic cortex, anterior insula and nucleus accumbens. These findings show that the vulnerability to develop compulsive cocaine seeking behavior stems from pre-existing structural or functional changes in two distinct cortico-striatal systems that underlie deficits in impulse control and goal-directed behavior.

## Introduction

Compulsion is a core component of addiction, defined by drug seeking and taking that persists despite personal harm (1, 2). Vulnerability to compulsion is hypothesized to result from an interaction between pre-existing individual differences in traits, environmental and experiential factors and long-term drug exposure (3-5). Thus, human stimulant addiction has been associated with novelty seeking (6), impulsivity (7-9) and impaired cognitive control (10) alongside alterations in both ventral and dorsal cortico-striatal circuits (9, 11, 12). However, the causal significance of such neurobehavioral vulnerability markers is unclear because they can be affected by drug exposure (7, 13-16).

Studies in rodents have been able to implicate causal neural substrates of impulsivity and novelty preference in vulnerability to compulsivity (3, 17-19). By contrast, novelty reactivity and sign-tracking (ST, the tendency to attribute incentive salience to Pavlovian cues) (20) confer an increased susceptibility to the acquisition of cocaine self-administration and sensitivity to the associative properties of cocaine-associated cues, respectively, rather than vulnerability to addiction (3, 18, 21). Nevertheless, none of these studies has acknowledged that compulsive drug use conflates phases of drug ‘foraging’ or anticipation, which actually occupies the majority of the time spent in waking activity of addicted individuals, prior to actual drug taking. Thus, they have principally investigated neurobehavioral circuitries implicated in compulsive stimulant drug-taking (22-24) following drug exposure thereby limiting our understanding of the neuropsychological basis of vulnerability to the compulsive tendency to seek drugs.

Compulsive drug use in preclinical models has generally been assessed in terms of the resistance to punishment of drug self-administration (25). Such self-administration behaviors exemplify instrumental (goal-directed) responding governed by the principles of reinforcement learning. However, the propensity for compulsive drug seeking in terms of possible predisposing differences in reinforcement sensitivity and the balance between goal-directed and habitual control over behavior (26, 27), has not yet been compared in conjunction with Pavlovian or neurobehavioral vulnerability markers.

Consequently, we combined a multidimensional behavioral approach to vulnerability with magnetic resonance imaging (MRI) to identify in drug naive male rats the structural and functional brain correlates of the later emergence of compulsive drug seeking. We therefore developed a novel behavioral procedure to investigate in individuals with a prolonged history of cue-controlled cocaine seeking (28) the individual vulnerability to persistent drug seeking in the face of punishment (**Fig. S1**).

## Materials and Methods

### Subjects and timeline of the experiment

As described in more detail in the **Supporting online materials (SOM)**, male rats underwent several MRI scans before being food restricted to 85% of their free feeding weight and screened for behavioral endophenotypes of vulnerability or resilience to cocaine addiction *(3, 29)*. As summarised in **Fig.S1**, rats were first screened for the trait of sign-tracking (ST) (20), then impulsivity *(18)*, reversal learning (15), followed by locomotor reactivity to novelty *(30)*. Individual differences in approach responses to conditioned stimuli (e.g., sign-or goal-tracking trait) were assessed in an autoshaping task *(3, 20)*. Impulsivity was measured in the 5-Choice Serial reaction Time task (5-CSRTT) (31); reinforcement learning and stickiness (32) were measured in a spatial reversal learning task *(15)*, while locomotor reactivity to novelty was assessed using four open fields and a video tracking system (ViewPoint Behavior Technology®, Lyon, France) (3). Upon completion of behavioral phenotyping, at PND 212 -244, rats were MRI scanned prior to intravenous catheter surgery, after which they were singly housed for the duration of the experiment. Rats were then trained to seek cocaine in the presence of the response-contingent drug-paired cue under a FI15(FR10:S) second order schedule of reinforcement, as previously described (33) and detailed in the **SOM**. After protracted cue-controlled cocaine seeking history, the tendency to persist in seeking cocaine despite adverse consequences was assessed as the resistance of drug seeking to mild electric foot shocks, as described in detail in the **SOM**. All experiments were carried out in accordance with the (U.K Animals) Scientific Procedures Act (1986) under the UK Home Office project licenses (PPL 70/7587 & PPL 70/8072) held by BJ and DB, respectively and were approved by the University of Cambridge Ethics Committee. The number of animals used during each stage of this longitudinal study are summarized in supplementary **Tables S1** and **S2**.

### Drugs

Cocaine hydrochloride (kindly supplied by the NIDA Drug Supply Programme to DB) was dissolved in sterile 0.9% saline. Drug doses are reported as the salt form.

### Magnetic resonance imaging

#### Imaging acquisition

High resolution MRI was performed on a 9.4 T horizontal bore MRI system (Bruker BioSpec 94/20 Bruker Ltd. Coventry UK). Images were acquired under anaesthesia (isoflurane using the manufacturer-supplied rat brain array coil with the rat in a prone position as described in detail in the **SOM**.

#### Voxel based morphometry

Voxel-based morphometry (VBM) was performed to assess morphological correlates of impulsivity, stickiness and compulsivity. An *unbiased* whole-brain analysis approach was adopted to capture brain-wide significant differences in grey matter, as described in the **SOM**. images were first manually reoriented to match the orientation of a reference template image using the bulk manual registration tool in the Statistical Parametric Mapping (SPM) 8 (Wellcome Trust Centre for Neuroimaging, University College London, UK) toolbox SPMMouse (SPMMouse, Wolfson Brain Imaging Centre, University of Cambridge (34)). After correspondence was achieved, structural images were bias corrected and segmented into three different tissue classes (grey, white and cerebrospinal fluid). Tissue class images were then rigidly co-registered to a reference template image. Non-linear registration was achieved through the use of the diffeomorphic anatomical through exponentiated algebra (DARTEL) procedure (35). Using the DARTEL warp fields, images were then warped and modulated to match the newly generated template images (**Fig. S2**). All images were manually checked for accurate registration and segmentation. Modulated grey matter maps were smoothed with an isotropic Gaussian kernel of 0.45 mm to promote normality of the data and mitigate imperfections in image registration. As detailed in the **SOM**, a general linear model was used for voxelwise analysis on the smoothed maps. Three independent models were created with block designs to assess main effects of impulsivity (HI, n = 12; LI, n = 12), compulsivity (HC, n = 7; LC, n = 7) and stickiness (HK, n = 14; LK, n = 14). Two independent linear regression models were created to assess compulsive cocaine seeking (n = 39) and cocaine seeking under no punishment (n = 39).

Main effects were assessed with contrasts based on Student’s *t*, with a restricted number of assessments made to avoid type I errors due to multiple comparisons: i) HI < LI; ii) HC < LC; ii) HC > LC; iv) HK < LK; v) compulsive cocaine seeking (negative correlation); vi) cocaine seeking under no punishment (negative correlation), among others (**Fig. S3**). These were selected based on our previous findings that high impulsivity is related to reduced grey matter volume in the ventral striatum (36), thinning of the insula cortex (37) and evidence of grey matter abnormalities in stimulant-dependent individuals (38, 39). To control for multiple comparisons across voxels, cluster statistics were used. A cluster-forming threshold of p < 0.005 was used to generate clusters which were then considered further where p < 0.05 (*uncorrected* cluster-level significance). Small clusters (smaller than a 0.5 mm sphere equivalent, c. 130 voxels) were ignored in further mitigation of type I errors.

#### Functional connectivity analysis

Prior to functional connectivity analysis, voxel dimensions in the header files for both the structural (MT-weighted) and functional images were scaled by a factor of 10 to facilitate processing with software designed for human brain images. After pre-processing (see **SOM** for more details) (40), images were first manually reoriented to match a reference template, as described above and processed as described in the **SOM**. Registration accuracy was manually checked for each image (**Fig. S4**). Temporal spikes were then removed (3dDespike) followed by motion correction (3dvolreg). Excess motion was calculated through relative framewise displacement (FWD) as described in the **SOM**. To assess any residual motion effects, average FWD was regressed against global connectivity (calculated as the average correlation value for each subjects’ correlation matrix), regional connectivity (average row-wise correlation value for each subjects’ correlation matrix), and edgewise connectivity (correlation between each region-to-region value for each subject). No connectivity measure was related to average FWD (**Fig. S5**). Signal to noise ratio (SNR) and temporal SNR (tSNR) was also calculated for each image, with SNR and tSNR well in line with previously published reports (40) (**Fig. S5**). Following pre-processing, region-to-region functional analysis was carried out. The first eigenvariate of the BOLD timeseries was extracted for each region of interest (**Fig. S6**) (fslmeants) and the Spearman’s rho correlation coefficient was calculated pairwise between each region of interest (ROI) as a measure of functional connectivity (**Fig. S3**). Correlation matrices were subsequently used to investigate the relationship between connectivity and behavior. All correlations between ROI BOLD signals were corrected for multiple comparisons with a false discovery rate set at q = 0.05 (41).

### Data and statistical analyses

Data are presented as means ± 1 SEM, box plots (medians ± 25% and min/max as whiskers) or individual datapoints, and analysed using STATISCA 10 (Statsoft, Palo Alto, USA) as described in detail in the **SOM**. When the assumptions of normal distribution or homogeneity of variance were significantly violated, the data were log-transformed. Behavioral data were subjected to repeated measures analysis of variance (ANOVA). Significant interactions were analysed further using the Newman Keuls *post hoc* test or hypothesis-driven planned comparisons wherever appropriate. For all analyses, significance was set at α = 0.05. Effect sizes are reported as partial eta squared values (ηp^2^).

Dimensional relationships between behavioral variables were analysed using Pearson’s correlation coefficient, r. Following dimension reduction, as detailed in the **SOM**, descriptive statistical analyses (Factorial analyses) were carried out on variables with inherent theoretical construct such as premature responses during LITI sessions in the 5-CSRTT *(31)* (**see Table S1 and S2**) using a principal component extraction method, with a maximum number of factors set at n-1 where n refers to the number of variables used in the analysis, with a minimum eigenvalue of 1 and varimax rotation.

## Results

Following acquisition of intravenous cocaine self-administration, rats were progressively trained to seek cocaine daily for long (15 min) periods of time under the control of the conditioned reinforcing properties of the response-produced drug-paired conditioned stimulus (CS) under a second-order schedule of reinforcement (SOR) (42). The compulsive nature of cocaine seeking was then measured individually as the tendency to persistent cocaine seeking, in a drug free state, in the face of response-contingent mild foot-shock punishment.

High-compulsive (HC), low compulsive (LC), and intermediate compulsive (IC) rats identified according to an unbiased cluster analysis of the punishment resistance of cocaine seeking (43), represented 18%, 41%, and 41% of the cohort, respectively (**Fig. 1A**). HC rats originated from 5 litters, while IC and LC rats originated from 9 and 11 separate litters, respectively. Of these, four HC, eleven IC and seven LC rats were siblings from two, four and three independent litters, respectively. The incidence of HC rats, of which the genetic contribution accounted for only a 14.5% increased risk (Bayes’ law), is similar to that previously reported for compulsive cocaine taking (3, 44) and the prevalence of addiction in human cocaine users (45). HC rats received more shocks than the other groups during the first 15 min drug-free period (**Fig. 1B**) and during the entire daily session (**Fig. 1C**). This reflected the tendency to persist in cocaine seeking, as HC rats maintained higher levels of responding on the active lever during the first drug-free seeking period (**Fig. 1D**) and throughout the entirety of each punished session than the other groups (**Fig. 1E**). Consequently, HC rats continued to obtain the maximum number of cocaine infusions available daily whereas NC rats decreased significantly their cocaine intake across punished sessions (**Fig. 1F**). The development of punishment-resistant cocaine seeking shown by HC rats was not due to differences in their acquisition of intravenous drug self-administration or drug seeking compared with LC rats (**Fig. S7**), nor was it attributable to a differential sensitivity to pain, as assessed using a hotplate test (**Fig. S8**) (3).

**Figure 1.**
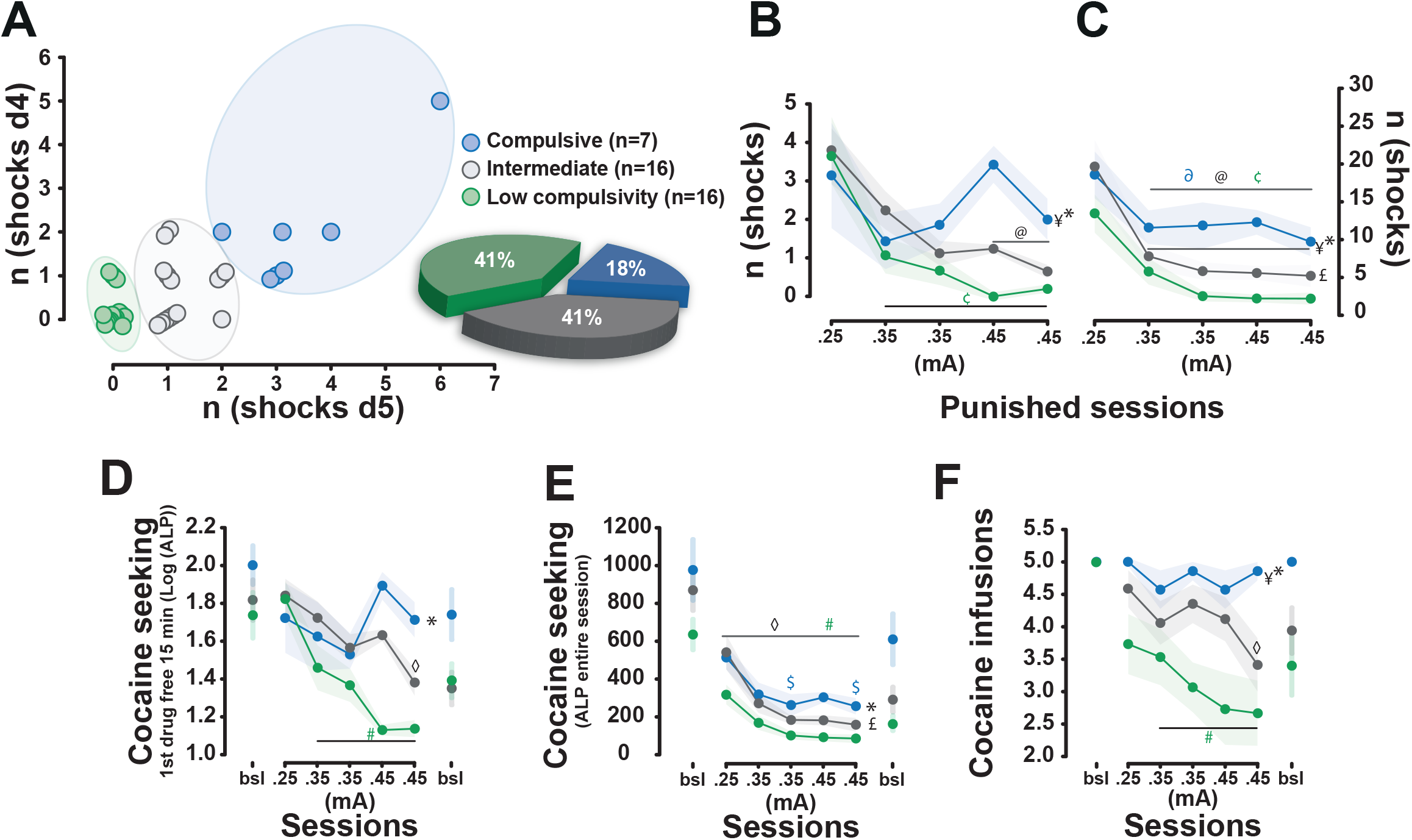
Emergence of a compulsive foraging phenotype in a subpopulation of rats with a long history of cue-controlled cocaine seeking. (**A**) Rats with a long history of cocaine seeking (n = 39) were identified by clustering as high compulsive (HC), intermediate (int) and low compulsivity rats (LC) based on the number of shocks they tolerated to pursue cocaine seeking over successive daily 15 min drug-free periods. (**B**) Under the threat of punishment, HC rats received more foot-shocks than Int and LC rats during the drug-free seeking periods **(B)** and the total session (**C**) [effect of compulsivity: F_2,36_ = 3.31, p = 0.047, ηp^2^ = 0.15 and F_2,36_ = 7.59, p = 0.0018, ηp^2^ = 0.29, respectively; compulsivity x session interaction: F_8,144_ = 2.15, p = 0.035, ηp^2^ = 0.11 and F_8,144_ < 1, respectively]. This willingness to persist in seeking cocaine in the face of punishment was reflected in higher levels of responding on the active lever in HC than LC rats during the first drug-free intervals [F_2,36_ = 3.29, p = 0.048, ηp^2^ = 0.15] (**D**) and the entire sessions (**E**) [F_2,36_ = 4.18, p = 0.023, ηp^2^ = 0.19]. (**F**) HC rats did not decrease their cocaine intake in the face of punishment as LC and Int rats [effect of compulsivity: F_2,36_ = 4.10, p = 0.025, ηp^2^ = 0.18]. n(shocks d4), n(shocks d5): number of shocks on the 4th and 5th punishment session; n(shocks), total number of shocks; mA, milliamperes; ALP, active lever presses; bsl, baseline session. Newman Keuls post-hoc tests, #, $, ◊: LC, HC, Int different from baseline, p < 0.05; ¢; ∂, @: LC; HC, Int different from d1, p < 0.05; *, ¥: HC different from LC and Int, p < 0.05; £: Int different from LC, p < 0.05.

Factorial analysis building on a dimension reduction strategy (see **SOM**) was used retrospectively on the putative behavioral markers of vulnerability (**Fig. S9**) to identify the detailed phenotype of drug-naïve rats later developing compulsive drug seeking (**Fig. 2**). These markers were premature responses in the 5-choice serial reaction time task (46), indexing impulsivity, the locomotor response to novelty (18), indexing novelty reactivity, conditioned approach to a food-related CS, indexing sign-tracking (3); reinforcement learning parameters (47) derived from a serial reversal learning task reflecting reward-based learning (α), reinforcement sensitivity (β) and ‘response stickiness’ (κ) (15). The latter parameter is the likelihood of the same response being repeated regardless of its reinforcing outcome and is thus a putative measure of value-free habitual responding (48). Importantly, this ‘stickiness’ is elevated in humans with stimulant use disorder (49) but it is unknown whether it is pre-disposing to addiction.

**Figure 2.**
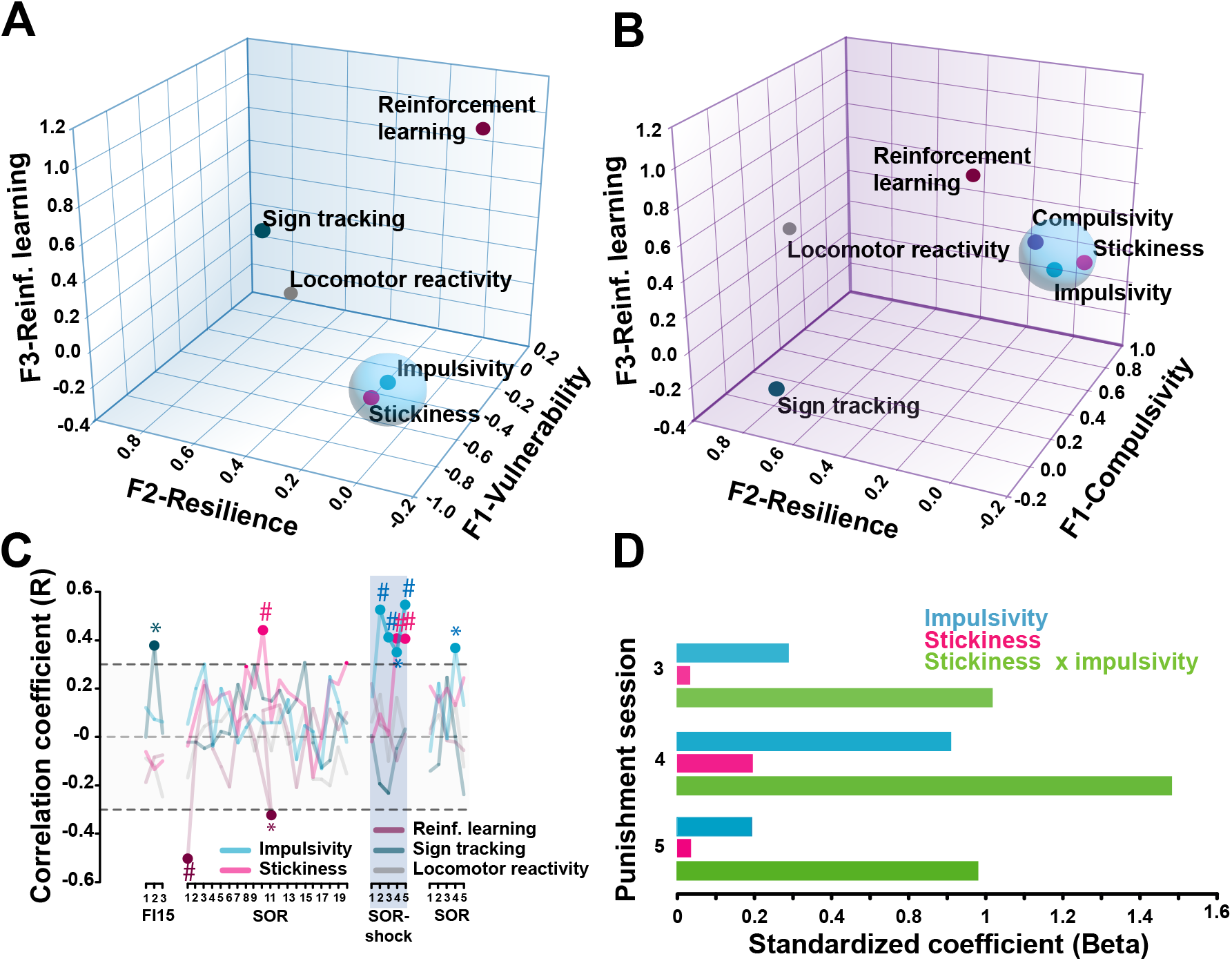
Impulsivity and stickiness interact to confer increased vulnerability to develop compulsive cocaine seeking. **(A)** A principal component analysis using locomotor reactivity to novelty, sign-tracking, the factor of reinforcement learning αβ, the stickiness factor κ, as well as impulsivity revealed three overarching factors accounting for more than 72.9% of the total variance (see **SM Table 1** for more details). Impulsivity and stickiness loaded on to factor 1, which represents vulnerability to compulsivity and was orthogonal to factor 2, which, by accounting for sign-tracking and locomotor reactivity-to-novelty, represents resilience to compulsion. Finally, reinforcement learning (αβ) loaded on the third, independent factor. **(B)** When compulsive cocaine seeking was added to the model, it loaded on the same factor as impulsivity and stickiness, while novelty reactivity and sign-tracking remained clustered on an orthogonal factorial capturing resilience, and reinforcement learning (αβ) was accounted for by a third independent factor. (**C**) Further dimensional analyses revealed that impulsivity and stickiness only correlated with cocaine seeking when under punishment, even though κ tended to correlate with cue-controlled cocaine seeking at baseline prior to the introduction of punishment. This suggests a relationship between stickiness and the tendency to engage in cue-controlled cocaine seeking. Grey area reflects the -.25 -+.25 marginal R value range. **(D)** Factorial regression analysis demonstrated that stickiness (κ) interacted with impulsivity in better predicting compulsive cocaine seeking on each of the last three days of punishment. *#: p < 0*.*01; *: p < 0*.*05*. FI15, fixed-interval 15 minutes; SOR, second order schedule of reinforcement.

Impulsivity and stickiness loaded on Factor 1 of the analysis (**Fig. 2A, Table S3**) while sign-tracking and novelty reactivity loaded on Factor 2, independent of Factor 3 which accounted for α and β (see **SOM** for more details). This multidimensional behavioral structure shows that more than half the overall model variance is explained by Factor 1 (vulnerability to compulsivity) and Factor 2 (resilience to compulsivity). Further factorial analysis incorporating compulsive cocaine seeking into this model (**Fig. 2B, Table S4**) revealed a shared construct of impulsivity, stickiness and compulsive cocaine seeking, loading on Factor 1, whereas Factors 2 and 3 accounted for resilience and reinforcement learning parameters, as in the initial analysis.

Subsequently, impulsivity was revealed to correlate with cue-controlled cocaine seeking only when rats had learnt that persisting in responding resulted in punishment, i.e. from the second punished session onwards (**Fig. 2C**). By contrast, stickiness not only correlated with cocaine seeking under punishment, albeit less systematically and robustly, but was also marginally related to baseline levels of cue-controlled cocaine seeking (**Fig. 2C**). Further analysis revealed that the co-occurrence of the behavioral traits of impulsivity and stickiness results in an increased vulnerability to develop compulsive cocaine seeking, the product of their variance being systematically a better predictor of compulsivity than that of each alone (**Fig. 2D**).

We next sought to define the neural basis of the complex endophenotype predicting compulsive cocaine seeking and the extent to which it mapped onto recent structural and functional neuroimaging studies of the propensity to stimulant use disorder in humans (9, 12, 50). Voxel-based morphometry analysis of scans carried out after behavioral screening and before any cocaine self-administration (∼post-natal day 228; (**Fig. S1**) (**Fig. 3, S3**) revealed that, compared to LC, rats destined to compulsively seek cocaine (HC) had lower grey matter density in the infralimbic (IL) cortex, as revealed by a negative correlation between compulsive cocaine seeking and the IL cortex density accompanied by an asymmetric distribution of HC and LC rats in the lower and upper terciles of the population ranked on IL density (**Fig. 3A**).

**Figure 3.**
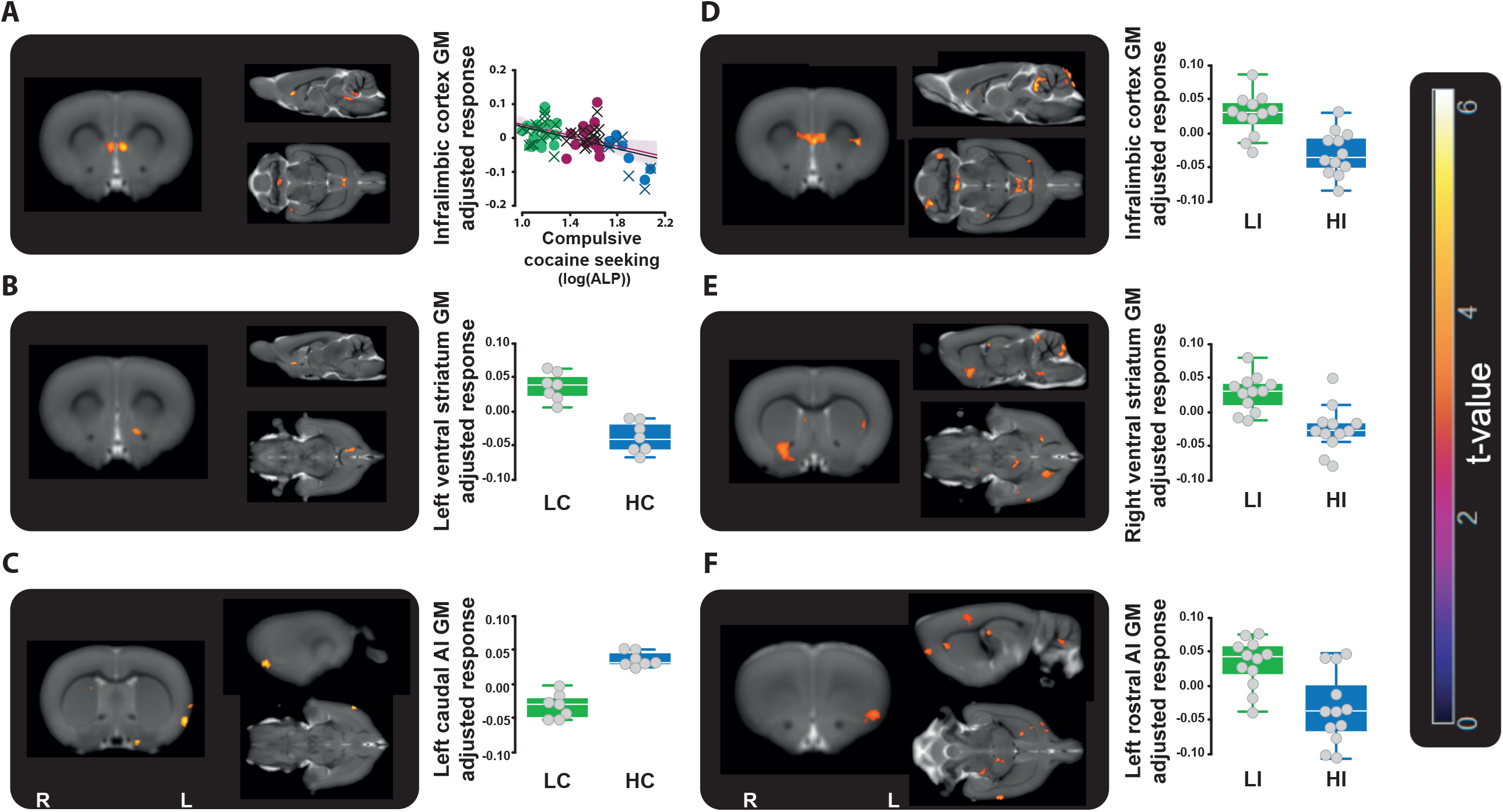
Grey matter alterations in the prefrontal cortex, the insula and the ventral striatum that underlie impulsivity predict compulsive cocaine seeking. Rats that will later compulsively seek cocaine (High-compulsive (HC) rats), showed prior to any drug exposure, lower grey matter density bilaterally in the infralimbic (IL) cortex **(A**), revealed by a negative correlation between compulsive cocaine seeking and the IL cortex density, (IL cortex, (left (.)) rho= -0.386, voxels = 13, cluster-level p = 0.013, (right (X)) rho= -0.356, voxels = 130, cluster-level p = 0.014), accompanied by a clear asymmetric distribution of HC and LC rats in the lower and upper terciles of the population ranked on the IL density [4 out of 7 HC rats were in the lower tercile as opposed to 3 LC which instead were predominantly represented in the upper tercile (10/16) which only contained 1 HC, Chi^2^: p = 0.0265] and a difference between groups in the left ventral striatum **(B)** [HC *versus* Low-compulsivity (LC) rats, voxels = 135, p = 0.003]. (**C**) Conversely, HC rats show higher grey matter density in the left caudal anterior insula (AI) (HC *versus* LC rats: voxels = 76, p = 0.019). This structural signature overlapped with that of High impulsive rats (HI) rats which, as compared to Low impulsive (LI) rats, showed lower grey matter density bilaterally in the IL cortex **(D)**, in the right ventral striatum **(E)** and the left rostral AI **(F)** [HI *versus* LI: voxels=176, p = 0.005, voxels = 397, p < 0.0001, and voxels = 154, p = 0.008, respectively]. Scatter plot and bar chart represent mean adjusted grey matter density from the significant cluster of interest. R = right hemisphere; L = left hemisphere. GM, grey matter; ALP, active lever presses

Rats destined to become compulsive also showed lower grey matter density in the left ventral striatum (**Fig. 3B**), but greater grey matter density in the left caudal anterior insula (AI) (**Fig. 3C**). This structural signature was specific to the tendency compulsively to seek cocaine as no such structural differences were observed in relation to baseline cue-controlled cocaine seeking (**Fig. S10**), and overlapped with the neural signature of impulsivity. Hence, as compared to Low Impulsive (LI) rats, High Impulsive (HI) rats showed lower grey matter density in the IL cortex (**Fig. 3D**), the right ventral striatum (**Fig. 3E**) and the left rostral AI (**Fig. 3F**), a neural profile consistent with earlier findings (36, 37). Together, these data suggest a rostro-caudal functional gradient in the AI mapping onto two different, yet interacting, behavioral manifestations of impulse control deficit, namely impulsivity and compulsivity, respectively (18, 37, 51-54).

The functional coupling signature (**Fig. 4A, Fig. S11C, D**) of compulsivity revealed a specific association with corticostriatal networks involved in goal-directed instrumental responding (55, 56). The coherence of coupling strength of the blood-oxygen-level-dependent (BOLD) response between either the prelimbic cortex (PrL) or the anterior cingulate cortex (ACC) with the posterior dorsomedial striatum (pDMS) was lower in HC compared to NC rats (**Fig. 4C**) but was not related to baseline cue-controlled cocaine seeking (**Fig. 4B**). Importantly, this hypofunctionality of the PrL→ pDMS and ACC→ pDMS that characterises compulsive cocaine seeking was also shown to underlie stickiness (**Fig. 4D**), but not impulsivity (**Fig. S11A, B**), a possible reflection of impaired goal-directed behavioral circuitry.

**Figure 4.**
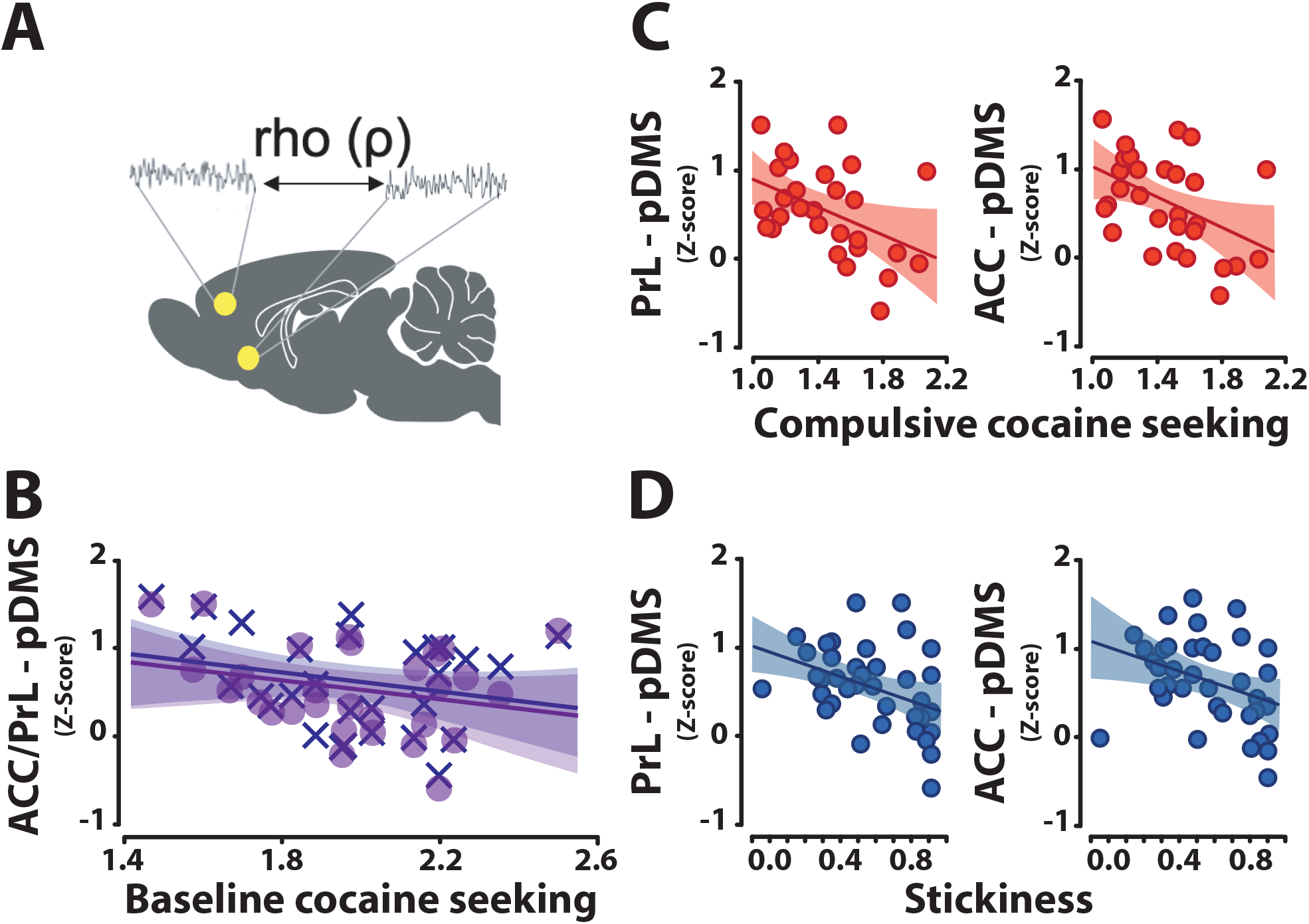
The functional hypoconnectivity of fronto-striatal regions that underlies stickiness predicts the vulnerability to develop compulsive cocaine seeking. (**A**) Several prefrontal-striatal regions of interest were assessed for their functional connectivity using Spearman’s Rho correlation coefficient. The functional connectivity between the anterior cingulate cortex (ACC) (X) and prelimbic cortex (PrL) (•) to the posterior dorsomedial striatum (pDMS), which was otherwise not significantly associated with cocaine seeking at baseline (**B**), was in drug naïve rats inversely proportional to the level of subsequent compulsive cocaine seeking [rho = - 0.441, p = 0.038 fdr-corrected and rho = -0.457, p = 0.030 fdr-corrected, respectively] (**C**). The hypofunctionality of these two systems in relation to compulsivity was similar to that observed in relation to stickiness (κ) prior to any drug exposure [rho = -0.463, p = 0.027, fdr-corrected and rho = -0.425, p = 0.048, fdr-corrected, respectively] (**D**). *(70)*

## Discussion

Our findings demonstrate the co-occurrence of high impulsivity and high stickiness traits increases the vulnerability to develop compulsive cocaine seeking in a large cohort of outbred male rats. These traits are related to structural and functional connectivity changes in dual neural systems that predict, prior to any drug exposure, the future transition to compulsive cocaine seeking. At the neural systems level, whereas impulsivity was accompanied by structural changes in the IL cortex, AI and ventral striatum, consistent with the recognised role of these regions in impulse control (36, 37, 57), the associated loss of control of cocaine intake (46, 58) or inflexible behaviors (59), stickiness, as indexed by κ, was associated with reduced connectivity strength between dorsomedial areas of the prefrontal cortex (PFC) (ACC/ PrL) and pDMS. This finding resonates with recent work showing increased stickiness in human stimulant users (49).

In humans, many studies have revealed structural and functional alterations of similar corticostriatal networks in SUD (9, 39, 60, 61), which might either be pre-existing vulnerability factors, but might also be the result of chronic drug exposure, or both, making it impossible to disambiguate causality. In contrast, our prospective longitudinal study has enabled resolution of this issue. Thus, we showed that the vulnerability to develop compulsive cocaine seeking is predicted by abnormalities in two distinct neural circuitries. One involved decreased functional connectivity between the ACC/PrL and pDMS, associated with increased behavioral stickiness. The second involved structural abnormalities in the IL cortex and ventral striatum, two key nodes of an impulsivity network associated with loss of top-down cognitive control (62).

Reduced connectivity between the medial PFC (mPFC) and pDMS, a region implicated in goal-directed behavior including drug seeking (63), is consonant with recent findings of a neurobehavioral endophenotype in this system in humans with stimulant use disorder and their unaffected relatives (50). It invites the hypothesis that pre-existing impairments in the goal-directed system lead to inflexible (‘sticky’), habit-prone behavior, which is expressed in compulsive drug seeking (27). This tendency to behavioral repetition independent of outcome may reflect the default operation of a value-free habit system that complements value-based reinforcement learning (48) and may underlie the profound perseverative tendencies of those addicted to cocaine, exacerbated by further failures of cognitive control (7, 64, 65). Pre-existing deficits in different circuits contributing to impulse control and flexible, goal-directed behavior may thus be joint precursors for the emergence of compulsive cocaine seeking.

The transition to severe cocaine use disorder (or dependence) has been reported to occur in only 15-20% of cocaine users (45), which is similar to the proportion of outbred rats shown here, that developed compulsive cocaine seeking, using an objective cluster analysis approach. This therefore is consistent with previous data showing that only 15-20% of male Sprague Dawley or Lister Hooded rats develop compulsive cocaine taking (3, 18). Furthermore, the combined statistical power of dimensional analyses and general linear models used here on a cohort of 39 individuals, together with between-subject comparisons on relatively smaller groups, confirmed the previously established relationship between impulsivity and compulsivity (18). This approach additionally allowed us to identify a novel interaction between impulsivity and stickiness in determining compulsive cocaine seeking vulnerability.

A limitation of the present study, however, is that it was confined to male rats in order to incorporate the results into a large existing dataset on addiction vulnerability mostly also confined to male rats (24, 66-69). Although severe cocaine use disorder is more prevalent in men than women, a necessary next step will be to undertake a longitudinal study in a large cohort of female rats to establish or otherwise the generalizability of these findings.

This identification of a complex neurobehavioral endophenotype for stimulant use disorder has implications for how we approach related psychiatric disorders, including addictions, as a necessary prelude for determining how the development of these fundamental behavioral systems is subject to polygenic and early experiential influences. This knowledge may ultimately help us appreciate how such putative genetic and environmental factors lead to the evidently profound individual differences in susceptibility to compulsive drug-seeking behavior that remain central to understanding and preventing substance use disorders.

The multi-modal approach we have adopted (with its neuroimaging and computational, as well as behavioral components) may also represent a more general strategy for modelling the etiology of other neuropsychiatric disorders.

## Acknowledgments

This work was funded by a programme grant from the Medical Research Council (MR/N02530X/1). J.A.J was supported by a Medical Research Council Doctoral Training Programme scholarship. B.J acknowledges funding from the AXA Research Fund, the National Health and Medical Research Council of Australia, and the Cambridge Newton Trust. P.Z was supported by a studentship from the Pinsent Darwin Trust in Cambridge. Cocaine hydrochloride was provided to D.B by the National Institute on Drug Abuse (NIDA) drug supply programme, Bethesda, USA. For the purpose of open access, the author has applied a Creative Commons Attribution (CC BY) licence to any Author Accepted Manuscript version arising.

## Author contributions

J.W.D, D.B, J.A.J, A.B.R, T.W.R, B.J.E and B.J contributed to the experimental design. J.A.J, L.H, B.J, C.V.S and P.Z contributed to behavioral screening. J.A.J, B.J, P.Z and S.J.S performed the imaging experiments. J.A.J, C.V.S and S.J.S analysed the structural imaging data. J.A.J analysed the functional imaging data. J.A.J, D.B, K.Z, P.Z, A.B.R and M.F performed the analysis of the behavioral data. D.B performed the intravenous catheter surgeries. A.B.R performed self-administration training with support from J.A.J and B.J. M.F and A.B.R analysed the cocaine self-administration data. M.F and D.B. designed and validated the compulsive drug seeking procedure. M.F. carried out the pain sensitivity experiment on an independent cohort. J.A.J, J.W.D, D.B, T.W.R and B.J.E wrote the manuscript. All authors contributed to editing the manuscript.

## Data and materials availability

All data needed to evaluate the conclusions in the paper can be found at: Zenodo. DOI: 10.5281/zenodo.7118542

