## Supplemental material for "Neurobehavioral precursors of compulsive cocaine-seeking in dual fronto-striatal circuits"

^#^ Joint last authors

† Corresponding author:

### Supporting Online materials and methods

#### Subjects

Male offspring of pregnant dams (Charles River, UK) were kept on a reverse light/dark cycle with red light on between 07:30am and 19:30pm and white light on between 19:30pm and 07:30am. Rats were MRI scanned on post-natal day (PND) 21 and weaned. Rats were MRI scanned again at PND 35 and 63, after which they were kept under food restriction to 85% of their free feeding weight during behavioral phenotyping. Upon completion of behavioral phenotyping, at PND 212 - 244, rats were MRI scanned prior to intravenous catheter surgery, after which they were singly housed for the duration of the experiment. All experiments were carried out in accordance with the (U.K Animals) Scientific Procedures Act (1986) under the UK Home Office project licenses (PPL 70/7587 & PPL 70/8072) held by BJ and DB, respectively and were approved by the University of Cambridge Ethics Committee. The number of animals used during each stage of this longitudinal study are summarized in supplementary **Tables S1** and **S2**.

#### Drugs

Cocaine hydrochloride (kindly supplied by the NIDA Drug Supply Programme to DB) was dissolved in sterile 0.9% saline. Drug doses are reported as the salt form.

#### Behavioral phenotyping

Beginning at PND 64, rats were screened for behavioral endophenotypes of vulnerability or resilience to cocaine addiction *(1, 2)*, as summarised in **Fig.S1** below. Rats were first screened for the trait of sign-tracking (ST) (*3*), then impulsivity *(4)*, reversal learning (*5*), followed by locomotor reactivity to novelty *(6)*. Individual differences in approach responses to conditioned stimuli (e.g., sign- or goal-tracking trait) were assessed in an autoshaping task *(1, 3)*. Impulsivity was measured in the 5-Choice Serial reaction Time task (5-CSRTT) (*7*); reinforcement learning and stickiness (*8*) were measured in a spatial reversal learning task *(5)*, while locomotor reactivity to novelty was assessed using four open fields and a video tracking system (ViewPoint Behavior Technology®, Lyon, France) (*1*).

##### Sign and Goal Tracking

The ST trait, which has been shown to be associated with the resilience to develop compulsive cocaine self-administration (*1*), was identified using a procedure based on our earlier work *(1)*. Two days prior to the beginning of autoshaping training for food (i.e., 45 mg food pellets, Bio-Serv, USA), food-restricted rats received 20 pellets in their home cage to minimise food neophobia. The experimental sessions were conducted for 6 consecutive days in 12 operant chambers (31.8 cm x 25.4 cm x 26.7 cm, Med Associates, St. Albans, USA) located within a ventilated sound-attenuating cubicle. Front and back panels of the test chambers were made of aluminium, while the right and left walls and the roof were transparent acrylic plastic with a stainless grid floor. A pellet dispenser was installed behind the front wall of each chamber, supplying food pellets to a food magazine located 2 cm above the grid floor. Each test chamber was illuminated by one 3-Watt light bulb during the experimental session. The first day of training was a habituation session in which rats were placed in the operant chamber (house light and fan turned on) for 30 min. The second day of training consisted of a single session of magazine training for the rats to learn where food pellets were delivered, 50 pellets being delivered under a 30 s-variable time (VT) schedule. Rats were then trained over 6 daily sessions to associate the presentation of a compound conditioned stimulus (CS) comprising a cue-light and a lever, deemed active in that it enabled the measurement of Pavlovian approach and contacts with the compound CS. Responding on that lever had no programmed consequences to the subsequent delivery of a food pellet (unconditioned stimulus: US). CS-US presentations were delivered according to a VT-90 schedule such that on termination of each variable time, the cue light was illuminated and the active lever was inserted for an 8 sec period. At the end of this period, the CS was turned off, the active lever was retracted, and one pellet was delivered in the food magazine. During the 8 sec interval, contacts with the compound CS measured as active lever depressions, were recorded as sign-tracking events while head entries into the food magazine were recorded as goal-tracking events. Inactive lever presses (uncoupled from the CS) were also recorded as an indicator of general activity. Individuals within the upper and lower quartiles of the population stratified on their level of sign-tracking (lever presses) over the last three sessions, were considered sign-trackers (ST) and goal-trackers (GT), respectively *(1)*.

##### Impulsivity

Impulsivity, as measured in the 5-CSRTT, is an endophenotype of vulnerability to develop compulsive behavior (*4, 9, 10*). The 5-CSRTT has been described previously (*7*). After one habituation session during which 5 pellets were placed in each of the five apertures and 5 pellets into the magazine in order to facilitate exploration of the chamber and nose-poking behavior, rats were trained to acquire the 5-CSRTT. Each session began with illumination of the house light and the delivery of a food pellet in the magazine. Collecting this pellet initiated the first trial. After a fixed inter-trial interval (ITI) of 5 sec, a light at the rear of one of the response apertures was briefly illuminated. Responses in this aperture within a limited hold period (5 sec) were reinforced by the delivery of a food pellet in the magazine (correct responses). Responses in a non-illuminated aperture were recorded as incorrect responses and were punished by a 5 sec time-out period. A failure to respond within the limited-hold period was deemed an omission and was punished by the house light being extinguished for 5 sec and no delivery of food reward. Additional responses in any aperture prior to food collection (perseverative responses) were recorded but not punished. Responses made in any aperture before the onset of the target stimulus or ‘premature’ responses, were punished by a 5 sec time-out period. Subjects were considered to have acquired the task (i.e., with a stimulus duration of 0.7 sec and an ITI of 5 sec) when their accuracy was greater than 80% and omissions were fewer than 20%. Rats then underwent three 60 min challenge sessions with a 7 sec long ITI (LITI), separated by two baseline 5 sec ITI sessions. Increasing the ITI results in a marked increase in premature responding and therefore facilitates the identification of inter-individual differences in impulsivity. HI and LI rats were selected in the upper and lower quartile of the population based on the average of the premature responses expressed during the last two LITI sessions (*9*).

##### Reversal learning

Spatial-discrimination reversal learning was assessed using twelve 5-choice operant chambers placed in ventilated, sound-attenuating cubicles (Med Associates, Georgia, VT), as previously described (*5*). Subjects were initially habituated to the apparatus over two days, with each session lasting 20 min. They were then trained to enter the magazine to trigger the illumination of a single stimulus light (left or right) and to respond in the illuminated aperture for food delivery under a fixed-ratio (FR) 1 schedule of reinforcement. Once rats had achieved 50 correct responses, food reward was successively delivered under FR2 and FR3 schedules to the same criterion within a 30 sec limited hold period. A failure to respond within the 30 sec period resulted in a 5 sec time-out. Once animals achieved criterion under a 5 sec inter-trial interval, they were tested for spatial discrimination followed the next day by a reversal of the stimulus-reward contingency. Rats were given 1 h to complete the discrimination task by achieving 9 correct trials across the previous 10 trials. During the 1-hour within-session reversal test, responses in the previously incorrect aperture were signalled as correct (and reinforced with a food pellet) whereas responses in the previously correct aperture were signalled as incorrect (and not reinforced with food) (**Fig. S12**).

A Q-learning model was used to simulate the reversal learning data as previously described (*5, 11*). The model, defined below, included three parameters: α, β, and κ. Model parameters were fitted to each animal’s reversal data individually and then compared using analysis of variance (ANOVA). The learning rate α determines how quickly the agent adjusts the expected value of a response following positive or negative feedback. High α values allow the agent to increase (or decrease) the expected Q-value for that response after the response is followed by a reward (or not). The inverse temperature parameter β regulates how much an agent explores by responding randomly or exploits what the agent learned about the responses to date. A low β value would lead an agent to rely on the expected Q-values of the responses and hence exploit what they have learnt about the responses already. A high β value would lead to exploration that under some circumstances may lead to higher rewarded outcomes. However, in the present reversal task, with deterministic outcomes, a high β value would result in fewer rewards. Finally, the choice autocorrelation parameter κ is a measure of stickiness, or how likely an agent will perform the same response again regardless of reward outcome. Values of κ close to 1 reflect an agent “sticking” to the previous response while κ values close to −1 reflect choice alternation. In the present task, high stickiness is not advantageous since it leads to a reduction in rewarded outcomes following contingency reversals.

The model used to generate α, β, and κ parameters for subsequent analyses is summarized as follows:

$$P(c_{t}=L|Q_{t}\left( L \right), Q_{t}\left( R \right), L_{t-1}, R_{t-1}) =\frac{exp(Q_{t}\left( L \right)/\beta+\kappa*L_{t-1})}{expexp \left( Q_{t}\left( L \right)/\beta+\kappa*L_{t-1} \right) + exp(Q_{t}\left( R \right)/\beta+\kappa*R_{t-1})}$$

whereby a larger κ results in greater probability of the choice c_t_ at trial t being the same as the choice c_t_ at trial t−1.

##### Locomotor reactivity to novelty

High locomotor reactivity to novelty predicts an increased tendency to acquire drug self-administration (*4*) but resilience to switch from controlled to compulsive drug intake (*1, 4, 12*). The locomotor reactivity to novelty test was conducted as previously described (*1*) in four infra-red illuminated open fields (50 x 50 x 50 cm) under a light intensity similar to that of the holding rooms (~ 550 Lux at the centre of the open field). Rats were placed in the open fields for two hours and their locomotor activity recorded throughout. Individuals whose total distance travelled was in the upper and lower quartiles of the population were considered High Responders (HR) and Low Responders (LR) rats, respectively (*1*).

##### Drug self-administration

###### Apparatus

Experiments were conducted using thirty-six standard operant conditioning chambers (Med Associates, St. Albans, VT, USA) enclosed within a sound-attenuating box containing a fan to eliminate background noise. Each chamber was equipped with two retractable levers (4 cm wide, 12 cm apart, and 8 cm from the grid floor), a cue light (2.5 W, 24 V) above each lever, and a white house light (2.5 W, 24 V) at the back of the chamber, in front of the levers. Silastic tubing shielded with a metal spring extended from each animal’s IV catheter to a liquid swivel (Stoelting, Wood Dale, IL, USA) mounted on an arm fixed outside of the operant chamber. Tygon tubing extended from the swivel to a Razel infusion pump (Semat Technical, Herts, UK) located adjacent to the external chamber. Lever presses, presentation of light stimuli, reward delivery, and data collection were controlled by a PC running MED-PC IV.

###### Intra-jugular surgery

Rats were implanted with a custom-made indwelling catheter into their right jugular vein under isoflurane anaesthesia (O_2_ carrier gas; 2 L/min; 5% for induction and 2-3% for maintenance and analgesia (Metacam, 1mg/kg, sc., Boehringer Ingelheim) as previously described (*13*). Following surgery, rats received daily oral treatment with the analgesic for three days and an antibiotic (Baytril, 10mg/kg, Bayer) for one week. Catheters were flushed with 0.1 ml of heparinized saline (50 U/ml, Wockhardt®) in sterile 0.9% NaCl every other day after surgery and then before and after each daily self-administration session.

###### Self-administration training

Rats were trained to acquire cocaine self-administration (0.25 mg/100µl/5.7sec/infusion) under continuous reinforcement over 4 daily, 2-hour sessions. Under this schedule, each active lever press resulted in drug infusion initiated concurrently with a 20 sec time out that included onset of a 20 s illumination of cue light positioned above the active lever (conditioned stimulus; CS), offset of the house light and retraction of both levers. Inactive lever pressing was recorded but had no scheduled consequence. Active and inactive lever assignment was counterbalanced, and a maximum of 30 infusions was available for this stage. Following these 4 daily sessions under continuous reinforcement, the daily schedule of reinforcement was changed to fixed intervals, increasing across daily training sessions from 1 min (fixed interval 1 min, FI1) to FI2, FI4, FI8, FI10 and eventually FI15 min (*14*). After three sessions under an FI15 schedule of reinforcement rats were trained to seek cocaine under the control of the drug-paired CS for 30 sessions under a FI15(FR10:S) second order schedule of reinforcement (SOR) (*15, 16*). Under this schedule of reinforcement, a CS is response-produced every tenth lever press while the drug is self-administered upon the tenth lever press after a 15 min interval has elapsed.

###### Compulsive cocaine seeking

For five daily sessions under SOR (SOR sessions 21-25), drug seeking behavior was punished when rats were actively engaged in responding for the drug, namely during the last 7 minutes of each 15-minute interval. Thus, during the last 7 min of each interval, mild electric foot-shocks (1 sec duration, 0.25-0.45 mA) dispensed by a scrambler (Med Associate St. Albans, USA) connected to the grid floor of the operant boxes were delivered on every 16^th^ lever press. Punishing responding only during the last 7 minutes of each interval offered the advantage that animals did not receive an electric foot-shock immediately after a drug infusion, which could result in counterconditioning and a decrease in the aversiveness of the shock (*17*). Each aversive stimulus was paired with a cue light located on the top middle region of the wall (independent from those paired with drug infusions) and rats received one shock every 16^th^ lever press to minimise the probability of co-occurrence of CSs and shocks and to avoid extinction of their drug seeking behavior. The shock intensity was increased from 0.25mA on the first punishment session to 0.35mA on sessions two and three to 0.45 mA on sessions four and five, as previously described (*18*).

Following the punished sessions, rats were re-exposed to five baseline SOR sessions to investigate their ability to recover their initial drug seeking behavior, or to display long-term behavioral adaptations to successive punishment sessions. A K-means cluster analysis *(19, 20)* was carried out on the number of shocks rats were willing to receive during the first, drug-free, interval of the last two punishment sessions, when shock intensity was 0.45mA and a significant and stable suppression of seeking and taking was observed in non-compulsive individuals, as shown in **Figure 1C** and **1E**. This approach identified non-overlapping populations stratified according to their propensity to persist in seeking cocaine in the face of adverse consequences.

##### Pain sensitivity measurement

To ensure that potential differences in resistance to punishment were not attributable to a differential sensitivity to a nociceptive (‘pain’) stimulus, high and low compulsive rats from an independent cohort were subjected to a hot plate test prior to punishment sessions *(1)*. Six hours following the 17^th^ SOR session, rats were placed on a hot plate (Ugo Basile, Gemonio, Italy), calibrated to remain at a stable temperature of 52ºC. The time elapsed before the appearance of pain-associated behaviors, including paw-licking and jumping, was measured and considered a direct indicator of pain threshold *(21)*. Rats were then immediately removed from the hot plate and returned to their home cage.

#### Magnetic resonance imaging

##### Imaging acquisition

High resolution MRI was performed on a 9.4 T horizontal bore MRI system (Bruker BioSpec 94/20 Bruker Ltd. Coventry UK). Images were acquired using the manufacturer-supplied rat brain array coil with the rat in a prone position. Structural images were obtained based on a three-dimensional multi-gradient echo sequence (TR/TE 25/2.4 ms with 6 echo images spaced by 2.1 ms, flip angle 6° with RF spoiling of 117°). The field of view was 30.72 × 25.6 × 20.48 mm^3^ with a matrix of 192 × 160 × 160 yielding isotropic resolution of 160 µm with a total scan time of 6 min 36 sec with zero-filling acceleration (25% in the readout direction; 20% in each phase encoding direction). Magnetisation transfer pulses (10 µT, 2 kHz off-resonance) were applied within each repetition to enhance grey-white matter contrast. Post-reconstruction, images from each echo were averaged after weighting each by its mean signal. A series of resting state fMRI (rs-fMRI) scans were carried out using a multi-echo planar imaging (EPI) sequence, with zero-filling (factor 1.33) used for readout and a partial-FT of 1.5 in the phase-encoding direction with an acceleration factor of 1.5. Scan parameters were: field of view = 28.8 x 21.6 mm^2^, matrix size = 64 x 48 x 40, echo time = 15 ms, repetition time = 1.832 s, 450 frames with a thickness of 0.5 mm. A total of 450 frames were collected, with sequential acquisition, and three echoes acquired for each frame. A total of 171 scan sessions occurred across 52 animals. Throughout all scans, rats were anaesthetised with isoflurane (1-2% in 1L /min O_2_: air 1:4). During the acquisition of rs-fMRI, isoflurane was reduced to 0.8% in 1L /min O_2_: air 1:4. Respiratory rate and pulse oximetry (SA instruments; Stony Brook, NY) were measured with anaesthetic dose rates adjusted to ensure readings remained within a physiological range. Body temperature was measured and regulated with a rectal probe and heated water system to 36-37°C.

##### Voxel based morphometry

Voxel-based morphometry (VBM) was performed to assess morphological correlates of impulsivity, stickiness and compulsivity. An *unbiased* whole-brain analysis approach was adopted to capture brain-wide significant differences in grey matter. Although ROI-based methods have potentially greater statistical power due to averaging over whole anatomical areas they assume that the underlying signal is consistent within that area. Voxel based methods do not rely on anatomical areas having a uniform signal throughout a given area and thus are better suited to detect effects that are more heterogeneous in nature. Nevertheless, in practice, with spatial smoothing included in the VBM procedure, and anatomical information used to interpret the findings, there is considerable overlap between these methods. Images were first manually reoriented (pitch, yaw, roll, x, y, z) to match the orientation of a reference template image using the bulk manual registration tool in the Statistical Parametric Mapping (SPM) 8 (Wellcome Trust Centre for Neuroimaging, University College London, UK) toolbox SPMMouse (SPMMouse, Wolfson Brain Imaging Centre, University of Cambridge (*22*)). After correspondence was achieved, structural images were bias corrected and segmented into three different tissue classes (grey, white and cerebrospinal fluid). Tissue class images were then rigidly co-registered to a reference template image. Non-linear registration was achieved through the use of the diffeomorphic anatomical through exponentiated algebra (DARTEL) procedure (*23*). Using the DARTEL warp fields, images were then warped and modulated to match the newly generated template images (**Figure S2**). All images were manually checked for accurate registration and segmentation. Modulated grey matter maps were smoothed with an isotropic Gaussian kernel of 0.45 mm to promote normality of the data and mitigate imperfections in image registration. A general linear model was used for voxelwise analysis on the smoothed maps. Three independent models were created with block designs to assess main effects of impulsivity (HI, n = 12; LI, n = 12), compulsivity (HC, n = 7; LC, n = 7) and stickiness (HK, n = 14; LK, n = 14). Two independent linear regression models were created to assess compulsive cocaine seeking (n = 39) and cocaine seeking under no punishment (n = 39).

Prior to any formal analysis, total intracranial volume (TIV) was correlated with each variable of interest. A correlation analysis between TIV and compulsivity, stickiness and cocaine seeking under no punishment showed no relationship with TIV (p > 0.3), whilst impulsivity was somewhat related to TIV (p = 0.065). A *two-way* t-test showed that HI rats showed lower TIV than LI animals (t = 2.271, p = 0.03), while no other behavioral measures showed group differences in TIV. As a result, TIV was not included in the impulsivity model (above). Although the removal of variables such as TIV can increase sensitivity by accounting for known variance in the data, if the confounding variable correlates with important modelling factors, effects that are accounted for by that variable will also be removed. Thus, although the evidence (p = 0.065) is not definitive for a linear relationship between TIV and impulsivity, there is stronger evidence (p < 0.05) of a group difference. Therefore, the benefits of including the confounding variable are unjustified by the resulting risk of losing sensitivity and that is why we did not include it in the model. Moreover, it is difficult to account for such a specific effect of the TIV variable in areas of the brain shown previously using a range of convergent techniques to relate to impulsivity (*24*). Nevertheless, TIV was included as a covariate in all other models to remove differences related to brain size among the rats. Main effects were assessed with contrasts based on Student’s *t,* with a restricted number of assessments made to avoid type I errors due to multiple comparisons: i) HI < LI; ii) HC < LC; ii) HC > LC; iv) HK < LK; v) compulsive cocaine seeking (negative correlation); vi) cocaine seeking under no punishment (negative correlation), among others (**Figure S3**). These were selected based on our previous findings that high impulsivity is related to reduced grey matter volume in the ventral striatum (*25*), thinning of the insula cortex (*9*) and evidence of grey matter abnormalities in stimulant-dependent individuals (*26, 27*). To control for multiple comparisons across voxels, cluster statistics were used. A cluster-forming threshold of p < 0.005 was used to generate clusters which were then considered further where p < 0.05 (*uncorrected* cluster-level significance). Small clusters (smaller than a 0.5 mm sphere equivalent, c. 130 voxels) were ignored in further mitigation of type I errors.

Laterality effects were included in our MRI analysis. As shown in **Figure 3**, HI and HC rats showed apparent differences in the laterality of grey matter changes in the nucleus accumbens. However, although there was some overlap in the membership of these two groups, these were not necessarily the same animals. Furthermore, we previously reported using combined VBM and ex-vivo protein analysis that the HI phenotype implicates dendritic spine abnormalities in *both* the left and right nucleus accumbens (*25*). However, in this study, presumed structural abnormalities in the right nucleus accumbens were evidently of an insufficient magnitude to cross the threshold for detection by MRI. We interpret these findings to suggest that impulsive and future compulsive behaviors are shaped by abnormalities in *both* the left and right nucleus accumbens but in an asymmetric way that may vary from one batch of animals to the next.

##### Functional connectivity analysis

Prior to functional connectivity analysis, voxel dimensions in the header files for both the structural (MT-weighted) and functional images were scaled by a factor of 10 to facilitate processing with software designed for human brain images. Pre-processing steps included 1) averaging of echo times, 2) removal of the first ten volumes, 3) brain extraction, 4) registration, 5) despiking, 6) motion correction, 7) spatial smoothing, and 8) bandpass filtering (*28*). Images were first manually reoriented to match a reference template, as described above. For each volume of the rs-fMRI time-series, the three echo times were averaged (3dcalc, Analysis of Functional Neuroimages, AFNI_16.2.07, [https://afni.nimh.nih.gov](https://afni.nimh.nih.gov/)) (*29*) and the first ten volumes discarded, to reach steady state (fslmaths). To improve registration, non-brain voxels were removed from both structural and functional images. Individually-registered grey, white, and cerebrospinal fluid tissue prior outputs from the VBM pre-processing procedure (above) were combined to obtain a mask excluding non-brain voxels. Following brain extraction, registration of functional images to a reference template was performed using a two-step registration approach. First, the brain-extracted functional image was registered linearly (6 DOF) to the brain-extracted structural (MT-weighted) image (FLIRT (FMRIB Linear Image Registration Tool (*30, 31*). Subsequently, the structural image was registered non-linearly to the template image (FNIRT (FMRIB Nonlinear Image Registration Tool (*31*)). Both linear transformation matrix and nonlinear warp-fields were combined to transform the functional image into template space to minimise interpolation effects. Registration accuracy was manually checked for each image (**Figure S4**). Temporal spikes were then removed (3dDespike) followed by motion correction (3dvolreg). Excess motion was calculated through relative framewise displacement (FWD) *via* the methods of Power et al., using a rat brain radius of 8 mm, as previously described (*32, 33*). All animals displayed minimal FWD and therefore none were removed from any formal statistical analysis. Motion parameter outputs from the motion correction procedure were subsequently regressed from the functional images (fsl_regfilt). Spatial smoothing was then applied using an isotropic 4.5 mm kernel (3dBlurInMask) and bandpass filtering was applied (0.01 - 0.1 HZ) (3dBandpass). To assess any residual motion effects, average FWD was regressed against global connectivity (calculated as the average correlation value for each subjects’ correlation matrix), regional connectivity (average row-wise correlation value for each subjects’ correlation matrix), and edgewise connectivity (correlation between each region-to-region value for each subject). No connectivity measure was related to average FWD (**Figure S5**). Signal to noise ratio (SNR) and temporal SNR (tSNR) was also calculated for each image, with SNR and tSNR well in line with previously published reports (*28*) (**Figure S5**). Following pre-processing, region-to-region functional analysis was carried out. The first eigenvariate of the BOLD timeseries was extracted for each region of interest (**Figure S6**) (fslmeants) and the Spearman’s rho correlation coefficient was calculated pairwise between each region of interest (ROI) as a measure of functional connectivity (**Figure S3**). Correlation matrices were subsequently used to investigate the relationship between connectivity and behavior. All correlations between ROI BOLD signals were corrected for multiple comparisons with a false discovery rate set at q = 0.05 (*34*).

#### Data and statistical analyses

Data are presented as means ± 1 SEM, box plots (medians ± 25% and min/max as whiskers) or individual datapoints, and analysed using STATISCA 10 (Statsoft, Palo Alto, USA). Assumptions for normal distribution, homogeneity of variance and sphericity were verified using the Shapiro–Wilk, Levene, and Mauchly sphericity tests, respectively. When these assumptions were significantly violated, the data were log-transformed. Behavioral data were subjected to repeated measures analysis of variance (ANOVA) with time (daily sessions or time bins) or blocks of sessions (baseline vs punished sessions vs post-punishment recovery for cocaine seeking) and levers (active vs inactive) or response (CS vs magazine approach) as within-subject factor, while groups (ST vs GT, HI vs LI, HK vs LK, HR vs LR and HC vs LC) were used as between-subject factors.

Significant interactions were analysed further using the Newman Keuls *post hoc* test or hypothesis-driven planned comparisons wherever appropriate. For all analyses, significance was set at α = 0.05. Effect sizes are reported as partial eta squared values (ηp^2^).

Dimensional relationships between behavioral variables were analysed using Pearson’s correlations. Descriptive statistical analyses (Factorial analyses) were carried out on variables with inherent theoretical construct such as premature responses during LITI sessions in the 5-CSRTT *(7)*, CS contacts in the autoshaping task *(1, 3)*, the computationally derived variable κ that represents stickiness in the reversal learning task (*35*) and the total distance travelled in the locomotor reactivity to novelty task *(36)* (**see Table S1 and S2**).

When several variables measured within the same procedure had potentially overlapping variance, they were subjected to dimension reduction. This was the case of the various variables and associated parameters measured in the reversal learning task, namely win-stay, lose-shift, α, β, κ, trials to criterion, perseverative errors, % correct trials and errors to criterion. Here, a first dimensional approach was carried out to identify variables that heavily correlated with one another (R > 0.5, p < 0.05 as a threshold value) so that only one representative variable was subsequently used in a following factorial analysis. In doing so, α, β, marginally related to one another (R = 0.42), but not to κ, were positively correlated with win-stay and lose-shift, respectively. α and β were both negatively correlated with the level of perseverative errors while the former was also negatively correlated with trials to criterion and errors to criterion. Thus, a factorial analysis was carried out on α, β, κ that yielded two factors, one that accounted for α, β (-0.84 each) and one for κ (-0.99), accounting for 47% and 33% of the total variance overall. The outcome of this analysis confirmed that κ can be used as a behavioral measure of stickiness while the individual projection on factor 1 (αβ) was used as the behavioral measure of reinforcement learning in subsequent analyses

All factorial analyses were carried out using a principal component extraction method, with a maximum number of factors set at n-1 where n refers to the number of variables used in the analysis, with a minimum eigenvalue of 1 and varimax rotation.

### Supporting Online Figures

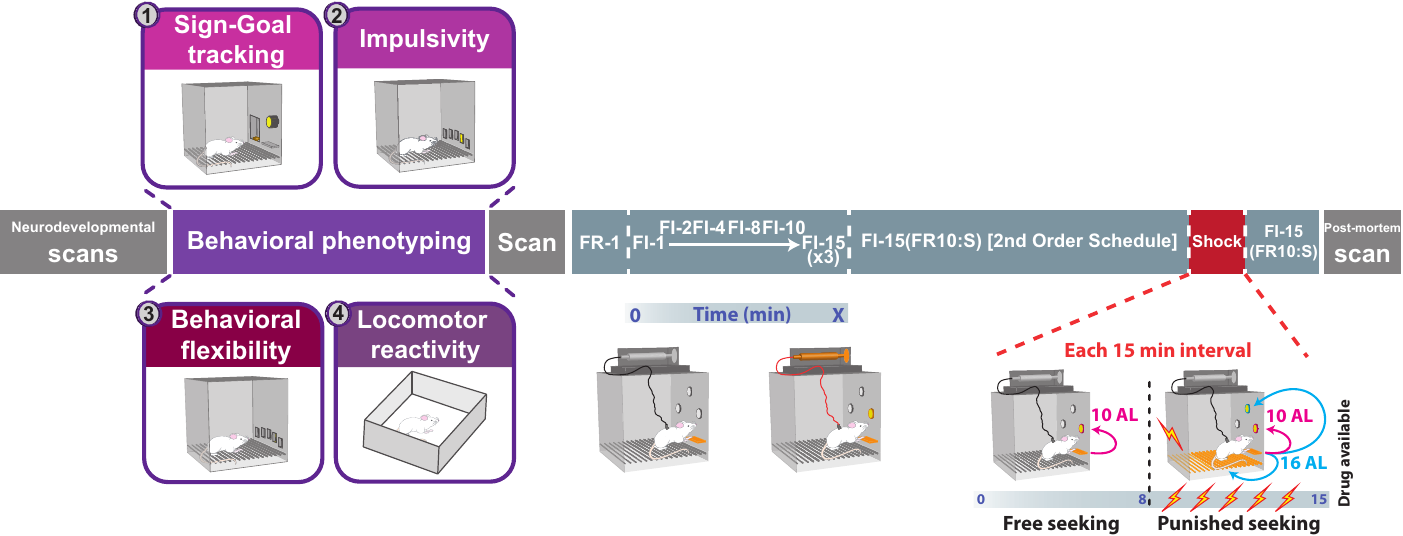

#### Figure S1. Experimental timeline.

Following three magnetic resonance imaging (MRI) scans over the course of the neurodevelopmental period, rats underwent behavioral phenotyping starting with their tendency to approach reward-predictive cues (sign-/goal-tracking), then their motor impulsivity (5-choice serial reaction time task), followed by their reinforcement learning performance and stickiness (spatial reversal learning), and their locomotor response to novelty. Following behavioral characterisation, rats underwent a further MRI scanning session prior to being implanted with intrajugular catheters. A week later rats were trained to acquire cocaine self-administration under continuous reinforcement (fixed ratio 1, FR1) over four daily sessions. Rats were then trained to seek cocaine under fixed interval schedules of reinforcement (FI) of increasing duration (from 1 minute – FI1- to 15 minutes -FI15) over the course of 8 daily sessions. Subsequently, rats were trained to seek cocaine under the control of the conditioned reinforcing properties of the drug-paired CS under a FI15(FR10:S) second order schedule of reinforcement (SOR) for 20 sessions (*62*). Compulsive cocaine seeking was then assessed by responding under punishment, in the form of mild foot shocks, for 5 sessions with increasing shock intensity, prior to re-baseline on SOR for 5 sessions.

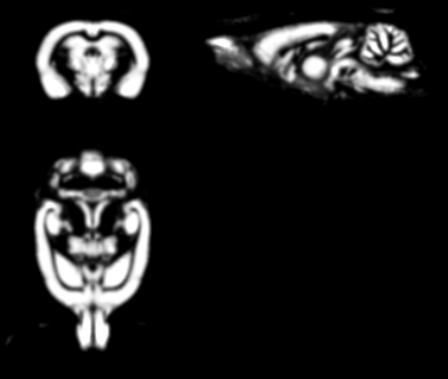

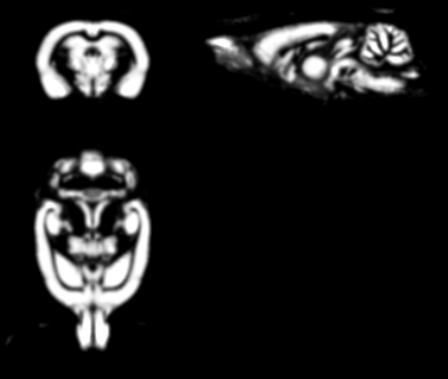

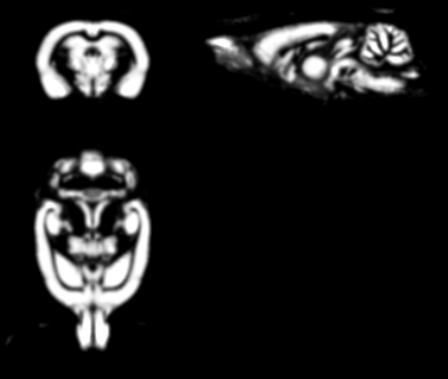

#### Figure S2. Representative smoothed normalised and warped grey matter volume map.

Voxel based morphometry was carried out on grey matter volume maps. All grey matter maps were checked for appropriate segmentation and normalisation prior to any formal analysis.

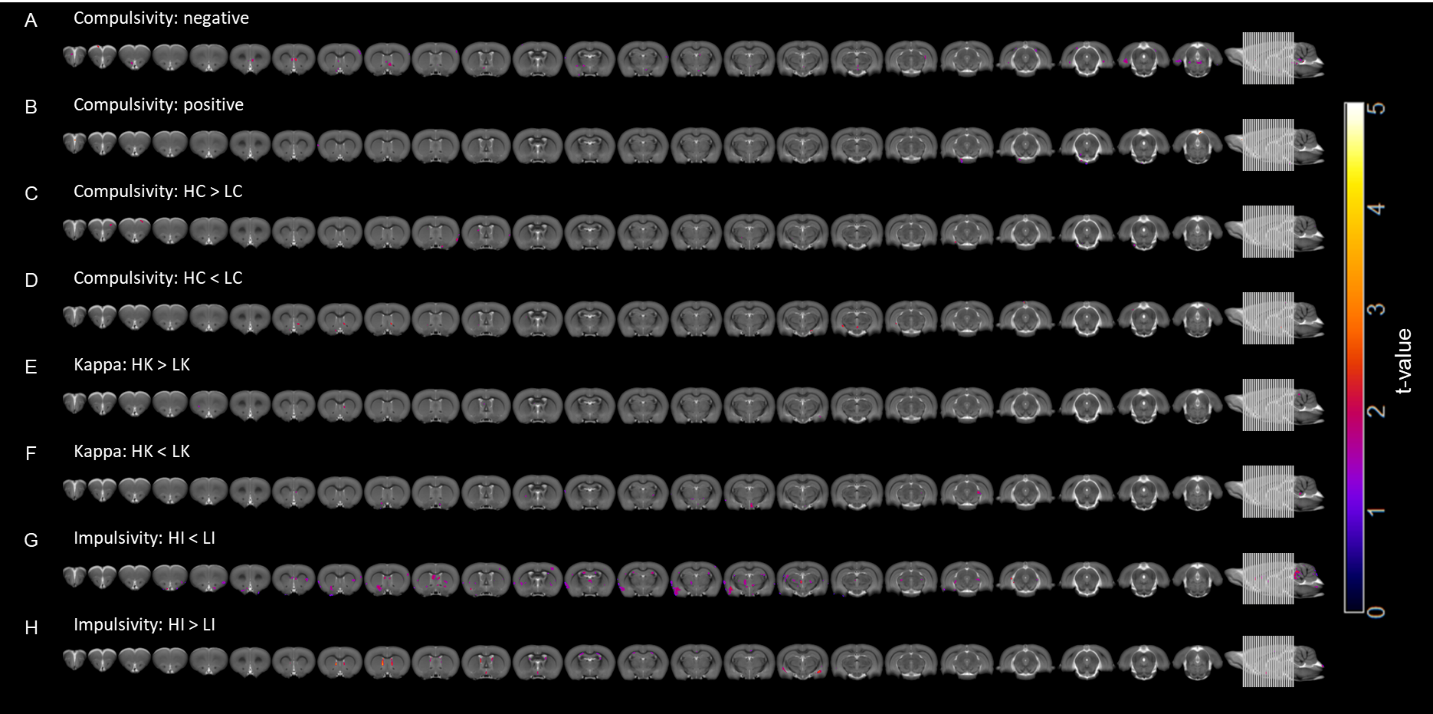

#### Figure S3. Voxel-based morphometric analysis of compulsive cocaine seeking, stickiness (kappa) and impulsivity.

**(A)** Regression (negative) analysis with compulsive cocaine seeking. **(B)** Regression (positive) analysis with compulsive seeking. **(C)** Compulsivity: high compulsive greater than low compulsive contrast. **(D)** Compulsivity: high compulsive less than low compulsive contrast. **(E)** Kappa: high kappa greater than low kappa contrast. **(F)** Kappa: high kappa less than low kappa contrast. **(G)** Impulsivity: high impulsivity less than low impulsivity contrast. **(H)** high impulsivity greater than low impulsivity contrast. All maps presented result from a chosen extent threshold of k = 15. Initial cluster-forming threshold p<0.005 (uncorrected).

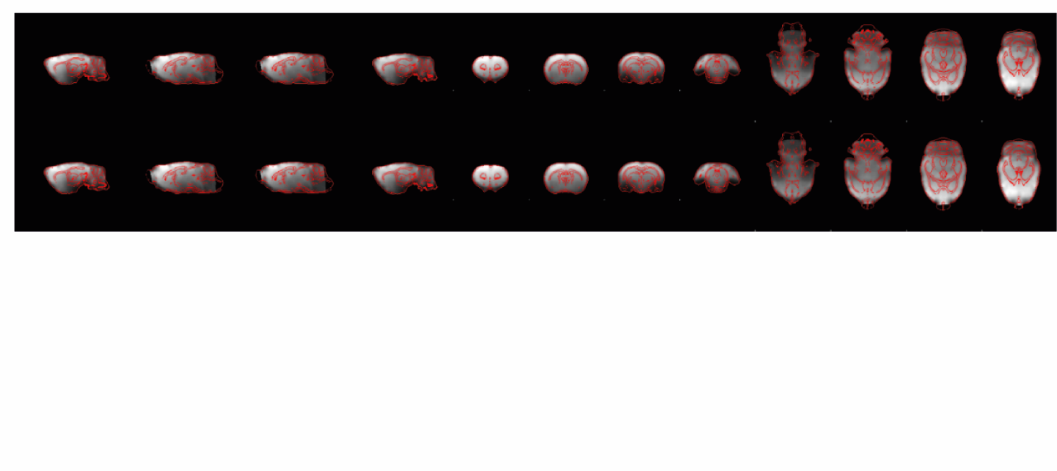

#### Figure S4. Representative normalization of resting-state functional MRI to study template.

Echo planar images were normalised to a study specific template using both linear and non-linear registration. Overlap between functional and structural template (red) is observed. All images were manually checked for appropriate registration efficacy. All scans were efficiently registered and none were excluded based on misalignment post-normalisation.

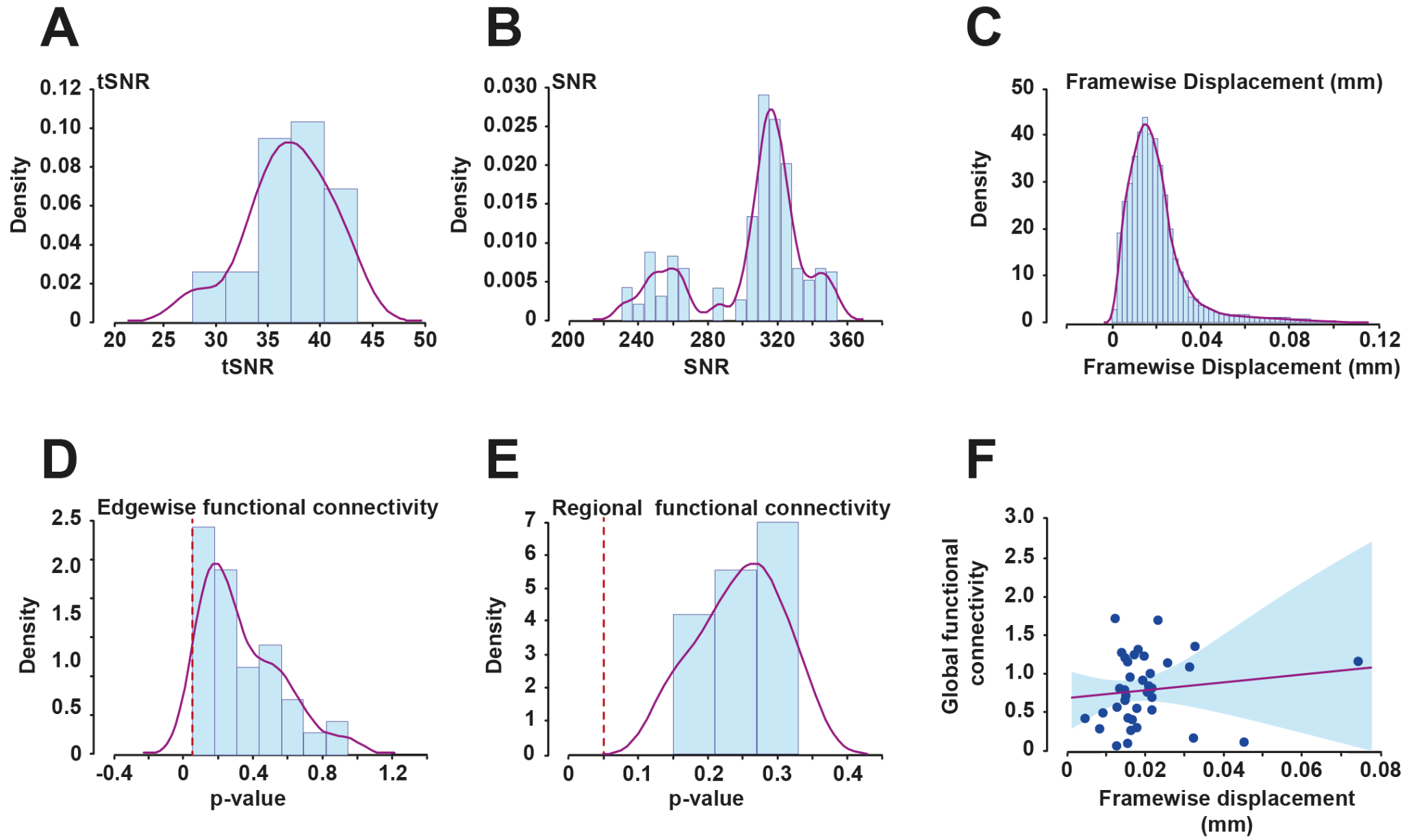

#### Figure S5. Quality control assessment of signal-to-noise, frame wise displacement and the evaluation of frame wise displacement on functional connectivity.

Temporal signal-to-noise (tSNR) was calculated by taking the average blood oxygenated level-dependent (BOLD) signal of all brain voxels, excluding non-brain voxels, across all volumes and dividing it by its standard deviation. Signal-to-noise (SNR) was calculated by subtracting the background noise signal from the global BOLD signal of several volumes across acquisition. Average tSNR was 36.95 (minimum: 27.74; maximum: 43.51; STD: 3.93), whilst average SNR was 305.44 (minimum: 230.50; maximum: 354.43; STD: 30.89). Framewise displacement (FWD) was used to estimate subject motion. Average FWD was 0.02mm, with maximum FWD 0.11mm. Several functional connectivity metrics were regressed against average FWD to evaluate the residual levels of motion on connectivity. Edgewise connectivity describes the region-to-region connectivity between a set of ROIs and showed no relationship with FWD (p > 0.05). Regional connectivity describes the average row-wise connectivity within the connectivity matrix and also showed no relationship with FWD (p > 0.05). Global functional connectivity, calculated as the average functional connectivity across each subjects’ correlation matrix, did not correlate with FWD (p > 0.05).

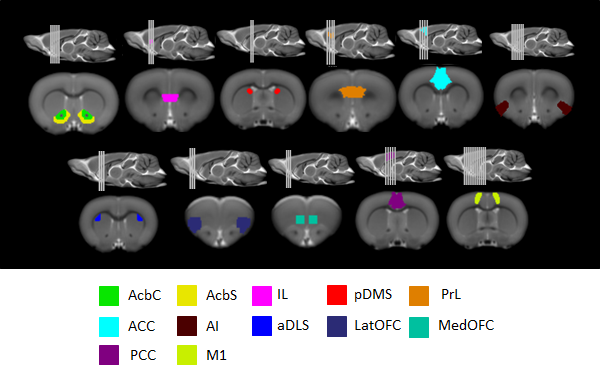

#### Figure S6. Regions-of-interest parcellation.

Manually drawn regions-of-interest were delineated based on Paxinos and Watson *(85)*. *Abbreviations*: AcbC, nucleus accumbens core; AcbS, nucleus accumbens shell; IL, infralimbic cortex; pDMS, posterior dorsomedial striatum; PrL, prelimbic cortex; ACC, anterior cingulate cortex; AI, anterior insula; aDLS, anterior dorsolateral striatum; LatOFC, lateral orbitofrontal cortex; MedOFC, medial orbitofrontal cortex; PCC, posterior cingulate cortex; M1, primary motor cortex.

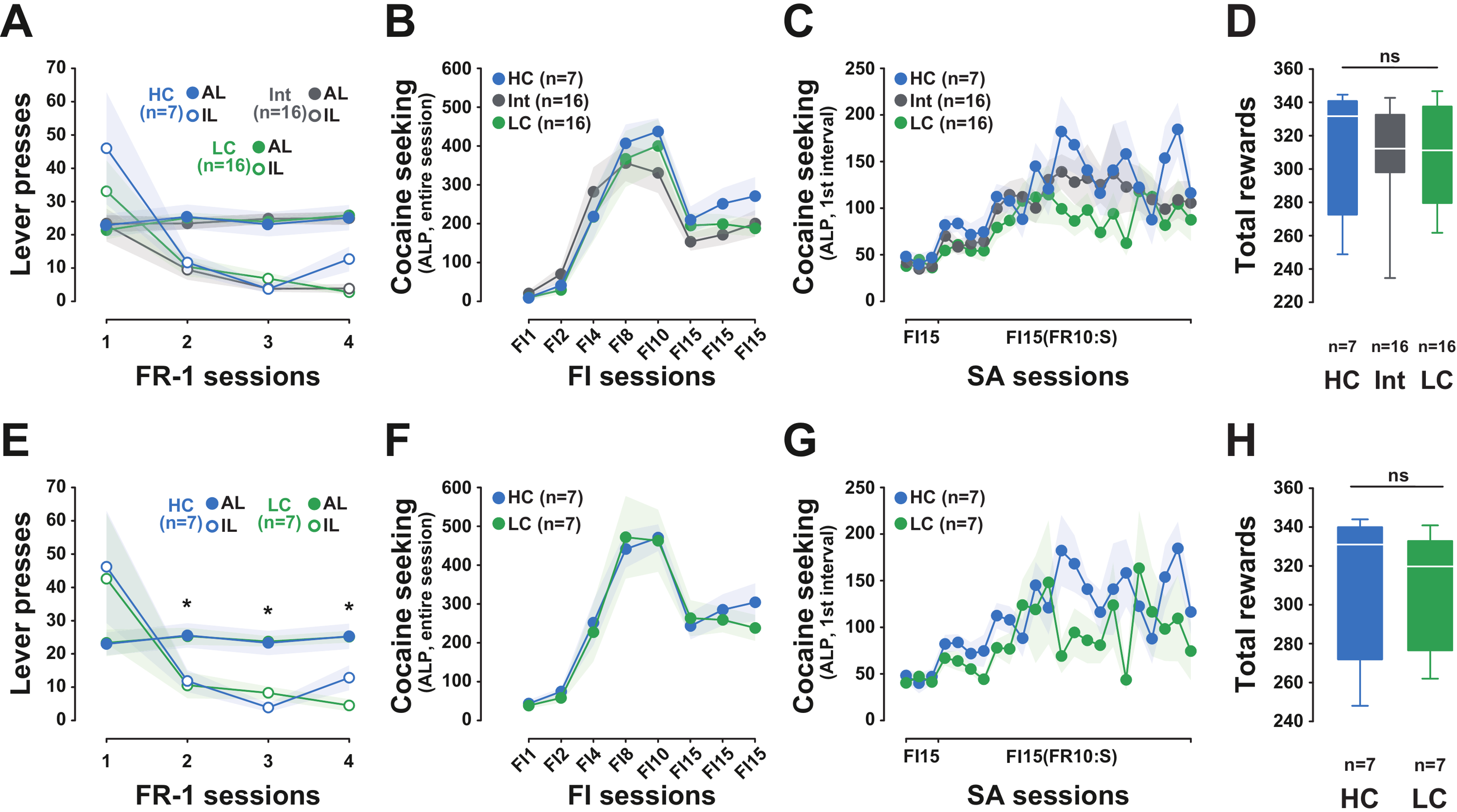
Figure S7. Individual vulnerability to persist in seeking cocaine in the face of punishment is not due to a difference in the rate of responding or to cocaine exposure prior to punishment.

**(A)** High-compulsive, Intermediate and Low-compulsivity rats (HC, Int and LC, respectively) showed a similar discrimination between the active lever (AL) and inactive lever (IL) and displayed similar levels of responding throughout the four cocaine self-administration acquisition sessions under continuous reinforcement [group x lever x session interaction: F_3,108_ = 1.473, p > 0.05]. **(B)** The three subpopulations did not differ in their acquisition of cocaine seeking under fixed interval schedules of reinforcement of increasing duration from 1 minute (FI1) to F15 minutes (FI15) [group x session interaction: F_7,252_ < 1]. **(C)** They did not differ in their sensitivity to the conditioned reinforcing properties of the cocaine-paired CS throughout their history of cue-controlled cocaine seeking under a FI15(FR10:S) second order schedule of reinforcement (SOR) [group x session interaction: F_22,792_ < 1]. **(D)** The three groups of rats received a similar number of cocaine infusions at the time compulsive drug seeking was assessed [main effect of group: F_1,36_ < 1]. **(E)** The seven HC, and seven LC rats selected for magnetic resonance imaging analysis showed a similar discrimination between the AL and IL and displayed similar levels of responding throughout all four cocaine self-administration acquisition sessions under continuous reinforcement [group x lever x session interaction: F_3,36_ < 1]. **(F)** HC rats did not differ from LC rats in their acquisition of cocaine seeking under fixed interval schedules of reinforcement of increasing duration from 1 minute (FI1) to F15 minutes (FI15) [group x session interaction: F_7,84_ < 1]. **(G)** HC and LC rats also did not differ in their sensitivity to the conditioned reinforcing properties of the cocaine-paired CS throughout their history of cue-controlled cocaine seeking under SOR [group x session interaction: F_22,264_ = 1.42, p > 0.05]. **(H)** Consequently, HC and LC rats received a similar number of cocaine infusions at the time when compulsive drug seeking was assessed [main effect of group: F_1,12_ < 1].

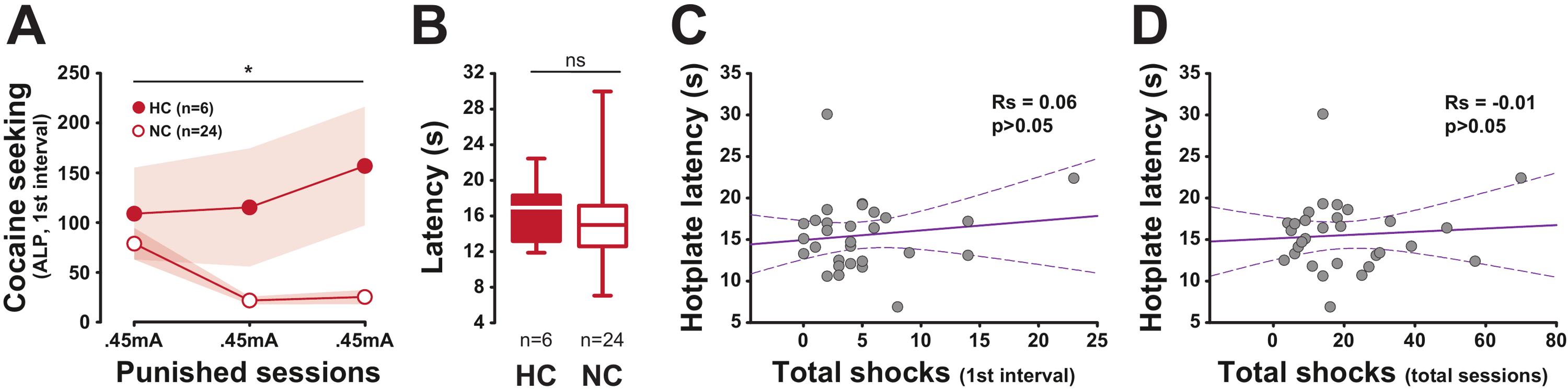

#### Figure S8. Individual vulnerability to persist in seeking cocaine in the face of punishment is not associated with differences in pain sensitivity.

(**A**) In an independent cohort of 30 rats, the introduction of punishment revealed a subpopulation of High-compulsive rats (HC, n = 6) that persisted in seeking cocaine despite the risk of receiving electric footshocks. In contrast, the majority of animals (Non-compulsive rats, NC, n = 24) decreased substantially their cocaine seeking responses [main effect of group: F_1,28_ = 10.78, p=0.002, ηp^2^=0.28; and session x group interaction: F_2,56_ = 3.93, p=0.02, ηp^2^=0.12]. (**B**) The difference in persistent cocaine seeking observed between HC and NC was not related to a difference in pain sensitivity as there were no differences in their pain threshold assessed by a hot-plate test prior to the punished sessions [main group effect: F_1,28_ < 1]. (**C**) Lack of relationship between pain sensitivity and the total number of shocks received prior to the first cocaine infusion. (**D**) Lack of relationship between pain sensitivity and the total number of shocks received over the 2-hour session. *: Newman Keuls post-hoc test, different from NC, p < 0.05.

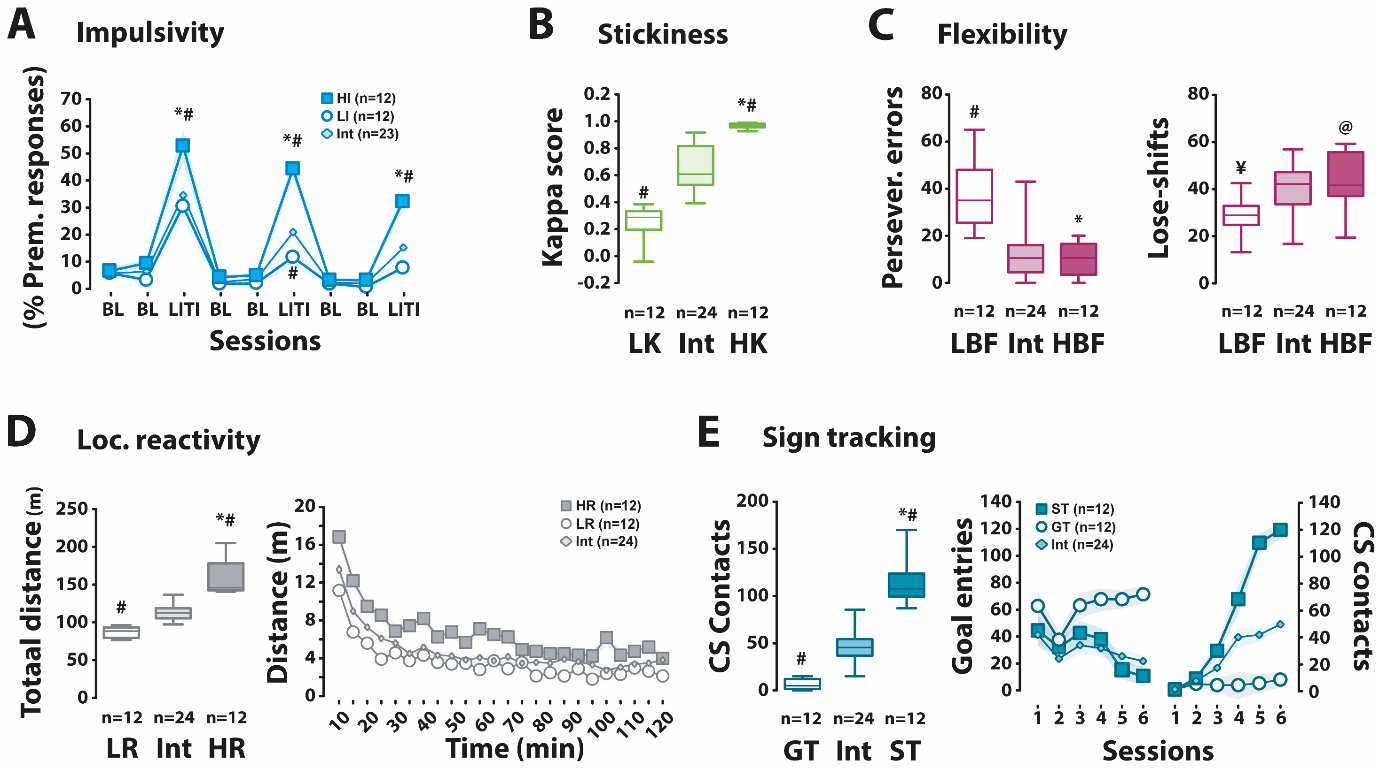

#### Figure S9. Multidimensional behavioral profile of heterogeneous rat populations.

Marked individual differences were revealed in the heterogeneous cohort of 48 Lister Hooded rats across several behavioral traits, including impulsivity as measured in the 5-CSRTT, reinforcement learning performance and stickiness, as measured in a spatial reversal learning task, sensation seeking, as measured by locomotor reactivity to novelty, and sign-tracking, as measured in an autoshaping task. **(A)** The introduction of longer inter-trial interval (LITI) in the 5-CSRTT increased the number of premature responses in all the groups (all ps < 0.001 vs baseline sessions), but much more so in highly impulsive rats (HI) as compared to intermediate (Int) and low impulsivity (LI) rats [main effect of group: F_2,44_ = 18.47, p < 0.0001, ηp^2^ = 0.45, session: F_8,352_ = 96.74, p < 0.0001, ηp^2^ = 0.69, and group x session interaction: F_16,352_ = 6.64, p < 0.0001, ηp^2^ = 0.23]. This differential effect of increasing the waiting time was specific to impulse control as attention, as measured as accuracy, was not affected by the LITI [group x session interaction: F_16,352_ < 1 even if it was overall slightly lower in HI rats (80-85% overall) than the other two groups (85-90% overall) [F_2,44_ = 6.23, p < 0.01, ηp^2^ = 0.22] (data not shown). Similarly, the increase in premature responses displayed by HI rats during LITIs was not due to an overall disengagement from the task as shown by the analysis the latency to collect the reinforcer [main effect of group: F_2,44_ < 1 and group x session interaction: F_16,352_ = 1.053, p = 0.40]. **(B)** In the reversal learning task, rats were stratified based on their kappa (κ) score, a proxy for stickiness. High κ rats (HK) displayed a higher κ score than intermediate and Low κ (LK) rats [main effect of group: F_2,45_ = 81.79, p < 0.0001, ηp^2^ = 0.78]. **(C)** Rats identified as showing high behavioral flexibility (HBF) based on their compound α/β score in the reversal learning task made fewer perseverative errors and more lose-shift responses than rats with low behavioral flexibility (LBF) [main effect of group: F_2,45_ = 21.62, p < 0.0001, ηp^2^ = 0.49, and F_2,45_ = 4.08, p < 0.03, ηp^2^ = 0.15, respectively]. **(D)** In the locomotor reactivity task, high responder (HR) rats reacted to a novel, inescapable environment much more than intermediate (Int) and low responder (LR) rats over a two-hour session [main effect of group: F_2,45_ = 86.58, p < 0.0001, ηp^2^ = 0.79, time: F_23,103 5_ = 80.83, p < 0.0001, ηp^2^ = 0.64, and group x time interaction: F_46,1035_ = 2.07, p < 0.0001, ηp^2^ = 0.08]. **(E)** In the autoshaping task, sign-tracking (ST) rats increased their contacts with the CS rather than the goal delivery location over the six daily training session, in contrast with intermediate (Int) rats that made equal contacts with the CS or the magazine, and goal-tracking (GT) rats that preferentially contacted the magazine, but not the CS [main effect of group x response x session interaction: F_10,225_ = 13.58, p < 0.0001, ηp^2^ = 0.38]. Thus, over the last three sessions, ST rats exhibited more conditioned approach to the CS than Int and GT rats [F_2,45_ = 119.81, p < 0.0001, ηp^2^=0.84].

Importantly, these traits were mostly independent of one another since HR and LR rats did not differ in their levels of impulsivity over the last two LITI sessions [F_1, 21_ < 1], sign-tracking [F_1,22_ = 1.63, p = 0.21], stickiness [F_1,22_ = 2.05, p = 0.166] or reinforcement learning performance, as assessed as the number of perseverative errors in the reversal learning task [F_1,22_ = 3.31, p = 0.082]. Similarly, HI and LI rats did not differ in their locomotor reactivity to novelty [F_1,22_ < 1], sign tracking [F_1,22_ < 1], or reinforcement learning performance [F_1,22_ = 1.22, p = .281].

However, HI rats showed higher stickiness (κ) than LI rats [0.73 ± 0.08 vs 0.46 ± 0.09, respectively, F_1,22_ = 4.677, p < 0.05, ηp^2^ = 0.17]. ST rats did not differ from GT rats in their levels of impulsivity [F_1,21_ < 1], stickiness [F_1,22_ < 1], or reinforcement learning performance [F_1,22_ = 1.65, p = 0.211]. However, ST rats displayed higher reactivity to novelty than GT rats [F_1,22_ = 6.74, p < 0.02, ηp^2^ = 0.16]. HBF rats did not differ from LBF rats in their level of reactivity to novelty [F_1,22_ < 1], stickiness [F_1,22_ < 1], impulsivity [F_1,22_ < 1] or sign-tracking [F_1,22_ = 3.20, p = 0.08]. HK rats did not differ from LK rats in their level of impulsivity [F_1,22_ = 3.67, p = 0.07], flexibility [F_1,22_ < 1], reactivity to novelty [F_1,22_ < 1] or sign-tracking [F_1,22_ = 1.02, p = 0.32].

\# and ¥: Newman Keuls post-hoc test, different from intermediate, p < 0.001 and p < 0.05, respectively, * and @: Newman Keuls post-hoc test, different from low trait group, p < 0.001 and p < 0.05, respectively.

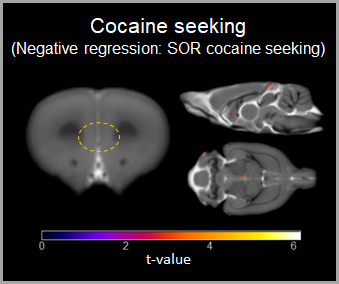

#### Figure S10. Cocaine seeking in the absence of punishment was not related to grey matter deficits in the infralimbic cortex or ventral striatum.

A voxel-based morphometric analysis evaluating the negative correlation between cocaine seeking under a second-order schedule (SOR) of reinforcement in the absence of punishment did not reveal significant associations with grey matter volume in the infralimbic cortex or ventral striatum, thereby confirming that the associations observed between seeking under the threat of punishment and alterations in these areas are indeed specific to compulsivity.

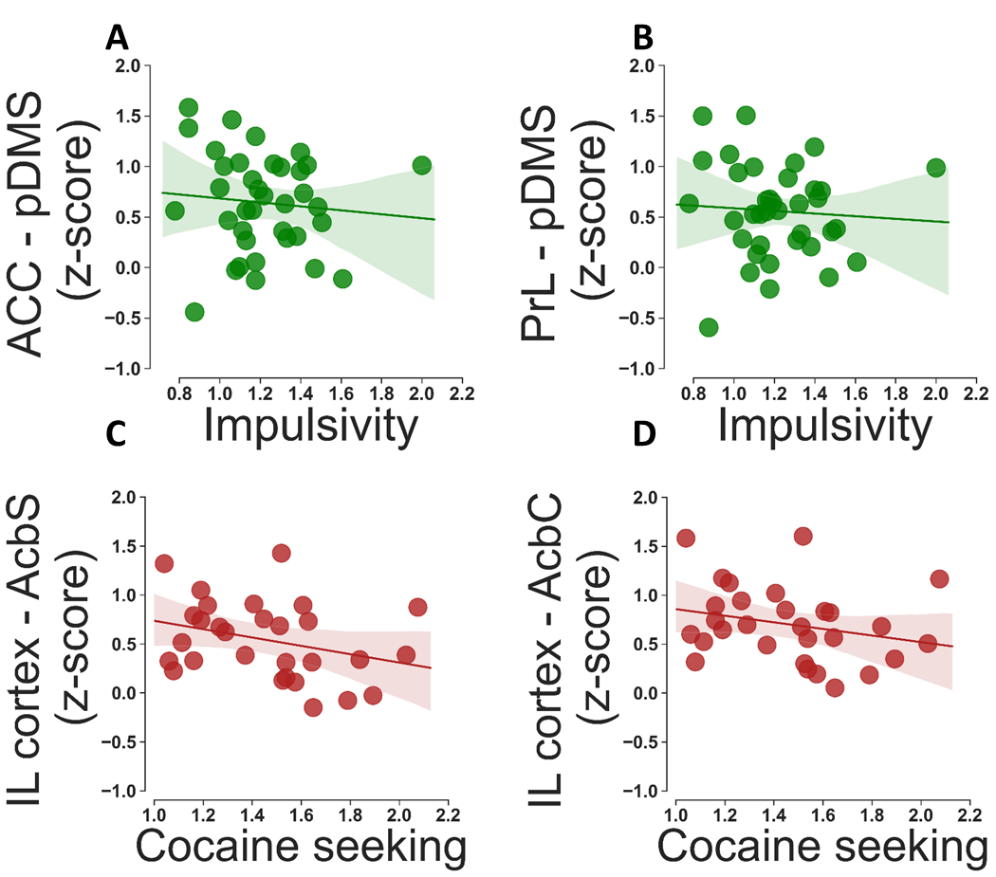

#### Fig. Figure S11. Functional connectivity strength between dorsal (ACC / PrL) or ventral (IL) prefrontal cortex and respectively pDMS and nucleus accumbens (AcbS / AcbC) does not correlate with impulsivity or compulsive cocaine seeking.

Relationships were analysed using Spearman’s correlation coefficient. Impulsivity was not significantly related to functional connectivity strength between (**A**) the ACC [rho = -0.148; p = 0.383] or (**B**) the PrL [rho = -0.132; p = 0.434]. Compulsive cocaine seeking was not significantly related to functional connectivity strength between (**C**) the AcbS [rho = -0.169; p = 0.316] or (**D**) the AcbC [rho = -0.141; p = 0.406].

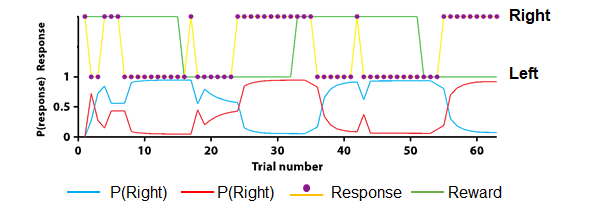

#### Figure S12. Representative reinforced learning modelling of serial deterministic reversal learning.

Model performance in the reversal learning task is illustrated using data from a representative animal. Responses made by the animal (left vs right, yellow trace and violet scatter points) are shown in the upper panel. The rewarded side (green trace, upper trace indicates that left responses were rewarded while lower trace indicates that right responses were rewarded) is also shown in the upper panel. Modelled probabilities of the same animal making a left or right response using the best fit values of α, β and κ are shown in the lower panel. Three reversals are shown. This animal required six trials to switch to the rewarded side after the first reversal, but quickly learned to switch more flexibly afterwards, taking only three trials to switch sides on the second and third reversals.

**Supporting Online Tables**

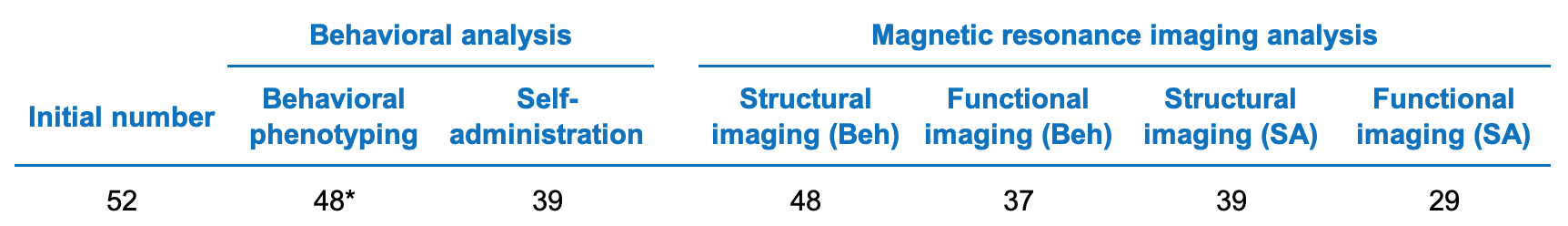

#### Table S1. Summary of the number of animals used in the study.

A cohort of n = 52 rats started the study (13 litters, each consisting of 4 rats). Due to technical difficulties encountered during MRI acquisition, three animals were excluded from the study. One further rat was euthanised due to malocclusion. During the cocaine self-administration procedure, nine animals were excluded due to illness or catheter patency issues. Due to sub-optimal quality assurance, eleven animals were excluded from the fMRI analysis. Rats destined to compulsively seek cocaine (n=7) were spread across five litters with one HC rat in each of three litters and two HC rats in each of a further two litters. Low compulsive rats (n=7) were each derived from separate litters. * indicates that one animal was removed from the analysis due to unstable behavior on the 5-choice serial reaction time task. Abbreviations: Beh, behavioral analysis (excludes compulsivity assessment); SA, self-administration procedure.

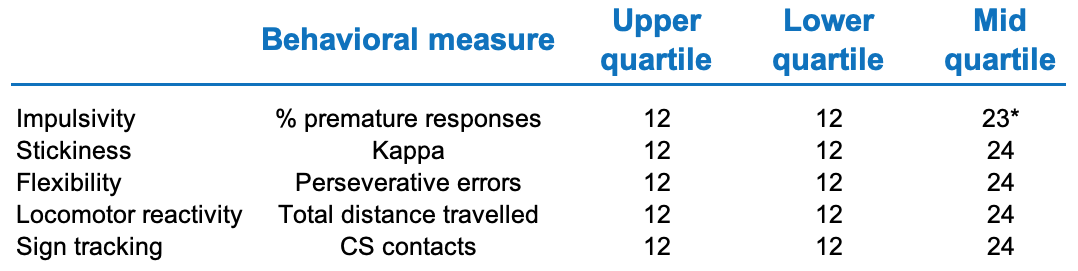

#### Table S2. Summary of the number of animals used for behavioral phenotyping.

Groups were based on the upper (75%) and lower (25%) quartiles of the ranked behavioral scores. * indicates that one animal was removed from the analysis due to unstable behavior on the 5-choice serial reaction time task.

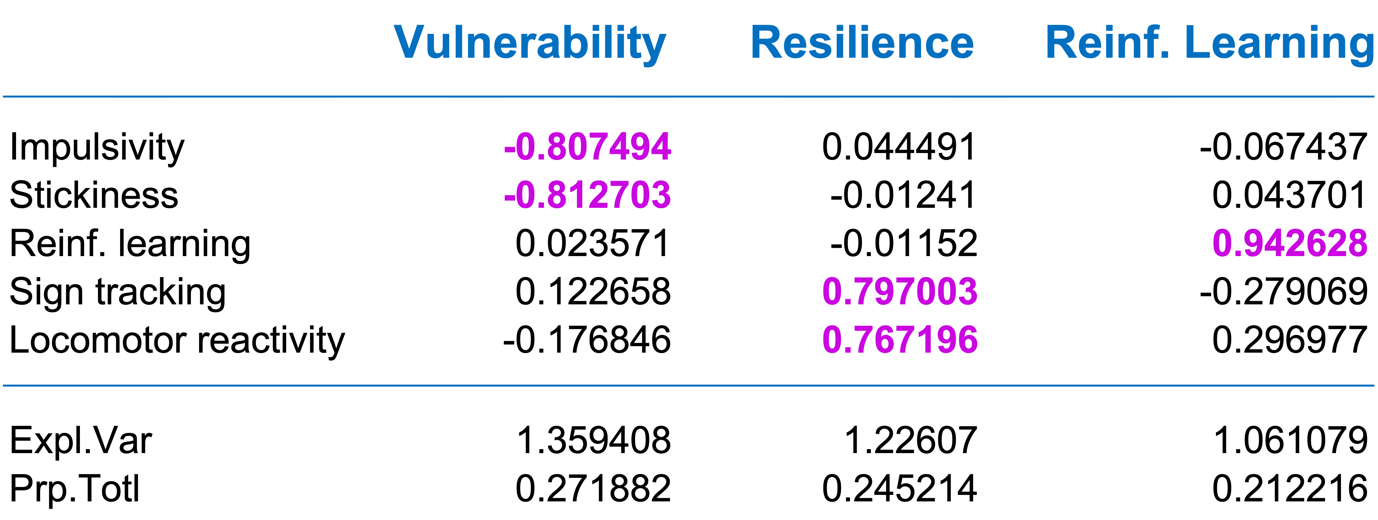

#### Table S3. Outcome of the Principal Component analysis on behavioral markers of vulnerability and resilience to compulsivity.

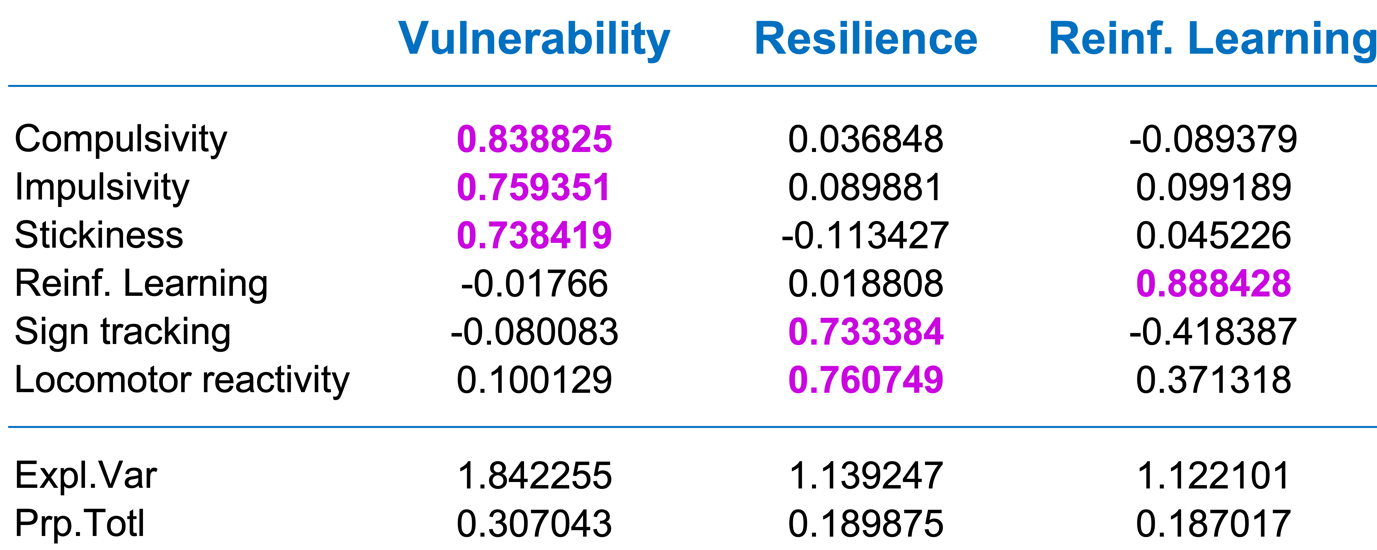

#### Table S4. Outcome of the Principal Component analysis on compulsive cocaine seeking and pre-existing behavioral markers of vulnerability and resilience.
